# Desync and connect: noninvasive MEG evidence of long-range coupling between locally desynchronized regions

**DOI:** 10.64898/2026.05.22.727161

**Authors:** Daria Kleeva, Alexei Ossadtchi

## Abstract

A prevailing assumption in systems neuroscience is that long-range phase coupling between cortical regions requires locally synchronized oscillatory activity within each interacting area. Here we ask whether distal phase coupling can be detected from MEG data when both interacting regions undergo concurrent event-related desynchronization (ERD). Using controlled simulations with progressively disrupted local phase coherence, we first establish that the Context-Dependent PSIICOS (CD-PSIICOS) framework – a projection-based connectivity estimator that separates power-dominated spatial leakage from interaction-specific cross-spectral structure – can recover distal coupling under increasing local phase dispersion, whereas estimators based on the imaginary part of the cross-spectrum fail at near-zero lags. We then analyzed recordings of a center-out reaching dataset from nine participants. A group decomposition of the coupling time courses identifies four long-range networks in alpha (8-13 Hz) and two in beta (15-25 Hz), each with its own time window. For each network we follow the local connections between its own sources as the power-dominated subspace is projected out, separating leakage-driven from leakage-resistant local coupling. In the earliest alpha network the leakage-resistant pairs stay at or below their pre-stimulus level at every rank while the power of the same sources falls, so this interhemispheric interaction may have no stable local counterpart. In the beta networks such a counterpart does survive, but the coupling it carries is an order of magnitude weaker than the long-range coupling of the same sources. Only in the latest alpha window, by which time power has returned to baseline, do local and long-range coupling rise together. Factorising the coupling into the amplitude of the single-trial cross-terms and the consistency of their phases separates the same cases: in the networks without a local counterpart, phase consistency increases inside the window while the amplitude of the cross-terms declines, whereas in the network whose local coupling rises with it, the increase reflects a growth of the cross-terms themselves rather than any improvement in their phase agreement. Long-range phase coupling can therefore persist, and in places strengthen, between regions undergoing macroscopic ERD, and noninvasive MEG can resolve this dissociation when leakage-aware estimation is employed.

## 1. Introduction

Neural synchronization is widely interpreted as a mechanism supporting functional integration across spatial scales. Phase-based measures such as coherence are used to quantify both local coordination within cortical regions and long-range coupling between distant areas. A prevailing assumption — most explicitly articulated in the communication-through-coherence (CTC) framework (Fries, 2005, 2015)— is that distal phase coupling is scaffolded by locally synchronized activity: stable intra-regional phase alignment creates temporal windows of excitability that render interareal transmission effective. Phase-based connectivity measures are defined only when an oscillation is present, so local desynchronization is naturally interpreted as eliminating the substrate for long-range phase coupling. However, a rhythm undetectable in noninvasive macroscopic recordings is not necessarily absent. Accordingly, throughout this paper we use “desynchronization” in the established sense of event-related desynchronization (ERD; (Pfurtscheller and Da Silva, 1999)), i.e. a macroscopic decrease of band-limited power.

Several lines of empirical evidence support this latter possibility and suggest that local and distal synchronization can dissociate. In sensorimotor paradigms, classical EEG work has demonstrated pronounced mu- and beta-band ERD over contralateral motor cortex during movement, coinciding with increased phase coherence between sensorimotor and frontal EEG sites rather than with its disintegration (Leocani et al., 1997; Classen et al., 1998). During visuospatial attention, alpha amplitude decreases contralateral to the attended location in early visual cortex, yet long-range alpha phase synchronization between visual and parietal regions is enhanced in precisely this hemisphere (Doesburg et al., 2009). More recent MEG work has consolidated the functional significance of such alpha power modulations: alpha lateralization causally biases spatial attention and sensory processing (Bagherzadeh et al., 2020), the cortical distribution of anticipatory alpha encodes non-spatial, object-based attention (de Vries et al., 2021), and the frequency and phase of alpha rhythms set the temporal windows within which visual input is integrated or segregated (Ronconi and Melcher, 2017; Gallina et al., 2024). These studies characterize alpha amplitude, its spatial distribution and its timing; the fate of long-range phase coupling in the very regions where alpha power decreases has received far less attention. Analogous dissociations have been reported during visual short-term memory retention, where posterior alpha desynchronization accompanies the increased frontal–posterior phase locking (Doesburg et al., 2010), and during object recognition, where alpha power decreases coincide with temporally and spatially specific increases in long-range alpha phase coupling (Freunberger et al., 2008). Importantly, these increases in long-range coupling are confined to task-relevant connections and time windows, arguing against explanations based on global arousal, nonspecific power changes, or volume conduction. More recently, Hindriks and Tewarie (2023) demonstrated mathematically and empirically that phase coupling networks and power correlation networks capture structurally distinct aspects of neural dynamics, with the dissociation driven by co-occurrent oscillatory bursts that contribute differently to each measure.

Several theoretical frameworks accommodate or predict such dissociations, yet none fully resolve the question we address. The gating-by-inhibition account (Jensen and Mazaheri, 2010) proposes that local alpha desynchronization reflects active disinhibition of task-relevant regions, compatible with concurrent long-range coordination, though subsequent work has questioned whether alpha reliably indexes local excitability or suppression (Foster and Awh, 2019; Jensen, 2024). Critically, this framework addresses whether a region can communicate but not how two simultaneously desynchronized regions achieve coherent interareal coordination.

CTC faces a parallel challenge: Schneider et al. (2021) provided intracranial evidence that long-range beta coherence between macaque premotor and parietal areas was intact without local spike-phase locking in the receiving area, proposing instead a Synaptic Source Mixing model in which interareal coherence arises as a consequence of communication through anatomical connections rather than as its cause. Related frameworks – Communication Through Resonance (Hahn et al., 2014), transient synchrony routing (Palmigiano et al., 2017), and coherence-through-communication (Vinck et al., 2023) – converge on the view that sustained local oscillatory synchrony is not a strict prerequisite for interareal phase coordination. At the computational level, Dahmen et al. (2022) showed that long-range coordinated activity can emerge in large-scale network models without strong local oscillatory order, with this result explained analytically by eigenvalue structure of heterogeneous connectivity matrices (Dahmen et al., 2019). At the cellular level, balanced excitation–inhibition decorrelates neuronal firing (Brunel, 2000; Renart et al., 2010), producing near-zero pairwise spiking correlations even when population-level field potentials exhibit oscillatory structure. Chimera state models (Kuramoto and Battogtokh, 2002; Abrams and Strogatz, 2004; Wang and Liu, 2020) further demonstrate how partial synchrony can maintain global integration while subsets of units remain desynchronized. Taken together, the information-via-desynchronization hypothesis (Hanslmayr et al., 2012; Klimesch et al., 2007) and the functional dissociation between local alpha amplitude and distal phase coupling (Palva and Palva, 2011) provide complementary perspectives suggesting that desynchronization may actively enhance local information coding rather than merely reflecting a breakdown of coordination.

Together, these empirical and theoretical lines of work suggest that distal phase coupling may, under certain conditions, persist independently of strong local synchronization. However, they leave a critical question unresolved. Existing studies either examine local desynchronization in one region paired with preserved synchronization in another, focus on amplitude-based measures without explicitly quantifying local phase dispersion, or interpret dissociations within task-specific functional frameworks. As a result, it remains unclear whether distal phase coupling can be maintained—or even dynamically enhanced—when *both* interacting regions undergo local desynchronization, whether such coupling specifically involves the very regions exhibiting macroscropic ERD (see Fig. 1) and whether such functional connectivity is implemented primarily via zero or close-to-zero phase coupling (Ossadtchi et al., 2018; Mehra et al., 2025).

**Figure 1:**
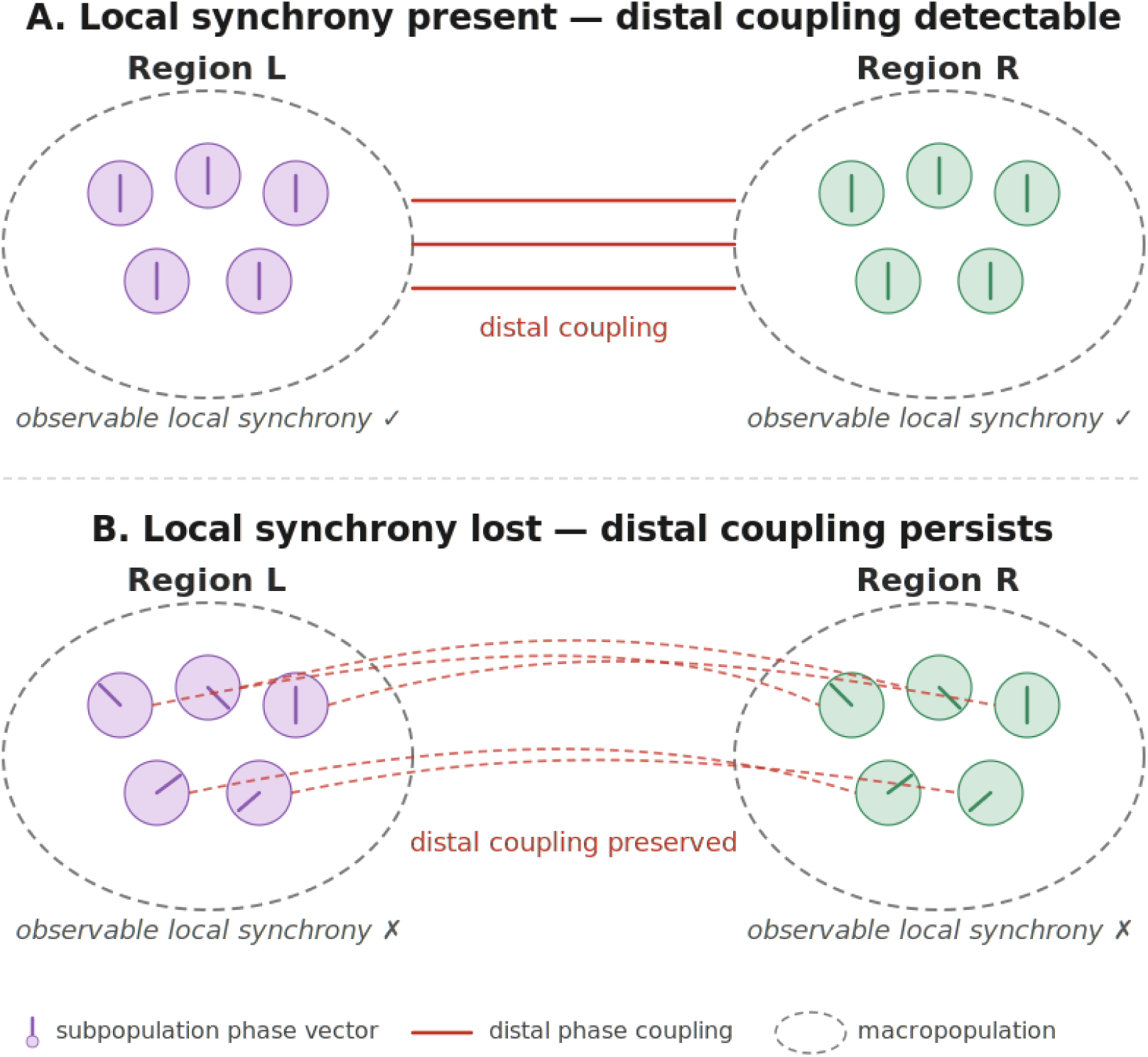
Schematic illustration of the dissociation between local and distal synchronization at the macropopulation level. Each dashed ellipse represents a cortical region of interest as resolved by MEG, containing multiple neural subpopulations (filled circles). Radial lines within each circle denote the instantaneous oscillatory phase of that subpopulation. (A) When all subpopulations within a region share a common phase (arrows aligned), the macropopulation exhibits observable local synchrony; distal phase coupling between regions L and R is readily detectable. (B) When subpopulation phases become dispersed (arrows misaligned), local synchrony at the macropopulation level is no longer observable. However, the corresponding subpopulations across regions maintain matched phase relationships, preserving distal coupling despite the absence of detectable local synchronization within either region.

Moreover, it remained technically challenging to test this question rigorously from noninvasive recordings, since conventional MEG/EEG connectivity estimators conflate local power contributions with interareal interactions especially in the instantaneous coupling scenario. Phase-based MEG measures have nevertheless proven capable of revealing functionally meaningful temporal coordination under non-strictly-periodic conditions, for instance in the attention-dependent cortical tracking of quasi-rhythmic visual input (Tabarelli et al., 2020). The present study addresses a related but distinct question: whether long-range brain–brain phase coupling can persist when local macroscopic oscillatory power is reduced. In the present study, powered by the novel and optimal set of tools (Kleeva and Ossadtchi, 2025; Altukhov et al., 2023) we directly address this question by systematically decoupling local and distal synchronization and explore their concurrent temporal dynamics.

First, using controlled simulations, we construct a generative model in which local phase coupling within two interacting regions is progressively disrupted while the phase relationship between the regions is held fixed by construction, and we test whether a leakage-aware estimator can recover the distal coupling from sensor-level MEG under these conditions. This is a detectability benchmark: it establishes that the regime of interest is recoverable when it is present, independently of task structure, signal power, or measurement confounds. Second, we test these predictions in empirical data using magnetoencephalography multi-subjects recordings from a center-out reaching paradigm, a task known to induce pronounced mu- and beta-band desynchronization in bilateral sensorimotor networks. Critically, rather than examining arbitrary inter-regional interactions, we focus on regions that exhibit strong local alpha and beta desynchronization and assess whether phase coupling between these same regions is preserved or dynamically strengthened over the course of the task. To estimate source-space functional connectivity while controlling for the confound between local power and interareal coupling, we employ the Context-Dependent PSIICOS (CD-PSIICOS) framework (Kleeva and Ossadtchi, 2025), a sensor cross-spectrum product-space projection-based method that separates the leakage subspace dominated by local power from the interaction-specific cross-spectral structure. We show that distal phase coupling does not merely survive local desynchronization in a set of isolated nodes, but exhibits a distinct and reproducible dynamic behavior illustrating transient interactions between regions that concurrently exhibit macroscopic ERD. Third, we ask whether these long-range interactions are accompanied by local phase structure in the same sources. Progressive removal of the power-dominated subspace distinguishes short-range coupling that decays together with the leakage from coupling that persists, and the level the latter attains indicates whether a locally synchronized subpopulation remains. Across the six networks the outcome ranges from the almost complete absence of a local counterpart, in an interhemispheric alpha network whose sources are strongly desynchronized, to the coexistence of local and long-range coupling in a late window in which power has already recovered.

## 2. Methods

### 2.1. Realistic simulations

At first we worked with simulated MEG data. We designed a set of simulations to test whether CD-PSIICOS can recover distal phase coupling from MEG-level data when within-region phase coherence, and hence macroscopic local synchrony and power, is progressively destroyed. The generative model isolates the two effects by construction: at fixed amplitude, within-region jitter disrupts the macroscopic coherent sum (and therefore sensor power), while a paired-jitter mechanism across regions keeps the inter-regional phase relationship intact.

Simulations were performed on the sample subject of the MNE dataset. A realistic three-compartment boundary element model with conductivities 0.3*/*0.006*/*0.3 S*/*m (scalp / skull / brain) was used to generate the forward operator, restricted to the *N_c_* = 204 planar gradiometers. A fine simulation source space (icosahedral spacing ico4, 5124 sources) was used to project source activity to sensors, and a coarser analysis source space (ico3, 1284 sources) was used for inverse modeling and connectivity analysis. The two source spaces use disjoint vertex sets, avoiding the inverse crime (Kaipio and Somersalo, 2005): each simulation vertex was mapped to the closest analysis vertex by Euclidean distance.

Simulated sources were current dipoles with fixed orientation normal to the cortical surface at each vertex of the simulation grid. In the analysis grid, each source was represented by the two leading spatial singular vectors of its free-orientation lead field, which for MEG span the quasi-tangential plane; all connectivity estimates aggregate over these two orientations (Section 2.3).

Two spatially compact ROIs were defined in the opposite hemispheres. Each ROI was grown as the *N_v_* nearest-neighbour vertices whose lead-field columns correlated *>* 0.3 with the seed lead-field, producing topographically consistent patches.

Each ROI was partitioned into *K* subpopulations of *N_v_/K* vertices; the subpopulation label of vertex *i* in the left ROI is denoted *s_L_*(*i*) ∈ {1*, …, K*}, and analogously *s_R_*(*i*) for the right ROI. In general the two assignments may differ; in the paired-jitter model used throughout this paper the *i*-th vertices of the two ROIs are assigned to the same subpopulation, *s_R_*(*i*) = *s_L_*(*i*) ≡ *s*(*i*), and we use *s*(*i*) below.

For each epoch *e* we drew a shared base phase φ^(e)^_base_ ∼ U[0, 2*π*) and a subpopulation-level jitter ε^(e)^_k_ ∼ N(0, σ^2^_local_), *k* = 1*, …, K*. Crucially, the same draw ε^(e)^_k_ was applied to the *k*-th subpopulation of *both* regions, so that within-region subpopulations drift relative to each other (destroying local synchrony) but the paired *L_k_*–*R_k_* phase difference is fixed. The phase of vertex *i* was therefore

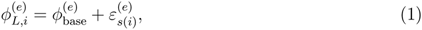

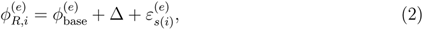

with an inter-regional phase lag Δ = *π/*20. Source time series were generated as unit-amplitude sinusoids at *f*_0_ = 10 Hz sampled at *f_s_* = 500 Hz,

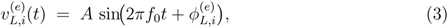

and analogously for *v^(e)^_R,i_(t)*. The amplitude *A* is *constant with respect to σ*_local_: the jitter does not modulate any vertex-level power; the observable power drop reported in the results arises entirely from subpopulation cancellation at the macroscopic level.

Two properties follow from the paired-jitter construction and are derived in Appendix A. Within-ROI coupling collapses: the mean cross-spectrum of two vertices of the same ROI carries a factor exp(−σ^2^_local_) unless they share a subpopulation, so in the *K* = *N_v_* regime used throughout (*K* = 30) it decays to zero as *σ*_local_ → ∞. Distal coupling instead saturates at a non-zero plateau: only the fraction 1*/K* of cross-region pairs shares its per-epoch jitter and the remainder decays exactly as within-region pairs do, so the cross-ROI coherent sum tends to *A*^2^*N* ^2^*/K*, and the limit *K* → ∞ corresponds to a fully unpaired model in which no distal coupling survives within-region desynchronization. Since both regions share a common per-epoch base phase, the simulated distal coupling is equivalent to a common drive with a fixed lag; the estimator quantifies the consistency of the inter-regional phase relation and does not distinguish direct interaction from shared input.

For every epoch the source activity was projected to the MEG sensors through the ico4 forward operator and additively contaminated with anatomically realistic background brain noise following the model of Ossadtchi et al. (2018). Specifically, *Q* = 1000 task-unrelated cortical sources were drawn at random (uniformly without replacement) from the simulation source space, with the two ROIs excluded from the pool, and re-drawn independently for every epoch. For each background source we generated a unit-variance white-noise time series and band-pass filtered it (4th-order zero-phase Butterworth) into four physiologically motivated bands — *α* (8–12 Hz), *β* (15–30 Hz), low-*γ* (30–50 Hz), and high-*γ* (50–70 Hz). The four band-passed traces were summed with weights inversely proportional to the band centre frequency (*w_b_* ∝ 1*/f*^-^*_b_*) to yield an approximately 1*/f* source-level spectrum, and then propagated to sensors through the ico4 lead field. The aggregate sensor-space brain noise was rescaled by a single global Frobenius factor so that

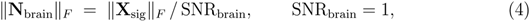

with the norms taken over the entire (epochs × channels × samples) batch. This batch-level Frobenius scaling, rather than per-trial normalisation, preserves the natural *σ*_local_-dependent decay of coherent sensor power that is the central observable of the desynchronization regime. Simulations were run for *T*_sim_ = 600 s per value of *σ*_local_, segmented into *N_e_* = 600 non-overlapping 1-second epochs.

The within-region phase dispersion was swept over *σ*_local_*/π* ∈ {0, 0.1, 0.2, 0.3*, …,* 2.0} (i.e. 0–2*π* radians). The chosen range covers the entire saturation curve from perfect local coherence to essentially uniform-random phase dispersion. For each *σ*_local_ the entire simulation was repeated independently for *N*_MC_ Monte-Carlo seeds; 95% confidence intervals in all subsequent plots are computed as ±1.96 SEM across these repetitions.

The subpopulation count *K* was treated as a free parameter and independently swept over *K* ∈ {2, 5, 10, 20, 30} to characterize how the preserved fraction of coherent cross-ROI pairs (1*/K*) shapes the distal plateau. The default regime used in the pipeline analyses corresponded to *K* = 30, which represents the most conservative case: absence of the genuine within-ROI local coupling.

### 2.2. Real data

The empirical validation was performed using an openly available MEG dataset acquired during a three-dimensional center-out reaching task (Yeom et al., 2023). The data were recorded with a 306-channel whole-head MEG system (VectorView, Elekta Neuromag Oy, Helsinki, Finland) comprising 102 magnetometers and 204 planar gradiometers at 102 sensor locations, inside a magnetically shielded room. Signals were sampled at 600.615 Hz with an online band-pass filter of 0.1–200 Hz. Participants performed visually guided reaching movements with the right arm toward one of four targets presented as three-dimensional anaglyph images. Each participant completed two recording sessions of 120 trials each (30 per target direction), yielding 18 recordings in total. The continuous recordings were processed with the spatiotemporal signal-space separation method (tSSS). Only pre-existing, fully anonymized data were used in the present study. All analyses in the present work were restricted to planar gradiometers.

The MEG data analyzed here were collected by Yeom et al. (2023). The experimental procedures of the original study were approved by the Institutional Review Board of Seoul National University Hospital (approval number 1105-095-363), all participants read and signed informed consent forms before the experiment and were informed that they could withdraw at any time, and the study was performed in accordance with the Declaration of Helsinki. The dataset is distributed under a Creative Commons Attribution 4.0 license. The present study is a secondary analysis of these openly available, fully de-identified recordings: no new data were collected and no participant-identifying information was accessed, so no additional ethical approval was required. All procedures were performed in compliance with relevant laws and institutional guidelines.

The analysis proceeds in six steps. (1) Sensor-level time–frequency analysis identifies the frequency bands and time windows of movement-related power suppression (Section 3.2.1). (2) Source-space power maps characterize the spatial distribution of that suppression and provide the weights of the CD-PSIICOS projector (Section 3.2.1). (3) CD-PSIICOS coupling is estimated for every source pair at a fixed projection rank, giving one coupling time course per pair per recording (Section 2.3). (4) These time courses, pooled over all recordings, are decomposed by temporal ICA into components that share a common time course and a fixed spatial weighting; components are retained on split-half reproducibility and on a surrogate test of their peak, and the network of a retained component is the set of pairs carrying its largest weights (Section 2.4; Sections 3.2.2 and 3.2.3). (5) Local pairs among the nodes of each retained network are followed across projection ranks and classified as leakage-resistant or leakage-driven; the accompanying tests quantify the consistency of these trends across recordings within the pre-selected pairs (Section 2.5; Sections 3.2.2 and 3.2.3). (6) The coupling of each network is decomposed into the amplitude of the single-trial leakage-free cross-terms and the alignment of their phases, and both are compared between the network’s own window and the pre-stimulus interval (Section 2.6; Section 3.2.4).

Continuous data were epoched from −1 to 5 s relative to target onset, and the trial-averaged evoked response was subtracted from each epoch to isolate induced (non-phase-locked) activity. All analyses reported below, including sensor-level time– frequency analysis, source power estimation, CD-PSIICOS coupling and the phase alignment analysis, were performed on these evoked-subtracted data. Sensor-level time–frequency representations were computed to identify frequency bands and time intervals associated with movement-related desynchronization. Time–frequency decomposition was performed using Morlet wavelet convolution (5–40 Hz in 1 Hz steps, number of cycles *n* = *f/*4, where *f* is the center frequency) on the induced part resampled to 100 Hz. Power estimates were baseline-corrected on a per-trial basis using a log-ratio transform. This normalization was applied for the purpose of identifying frequency bands and time windows exhibiting robust desynchronization.

All 18 recordings from the nine participants were analysed. A subset of eight recordings, from subjects 2, 5, 6 and 9, in which movement-related power suppression persists after the dominant CSP components are projected out, is characterised separately in Appendix E and used there as an illustration of the similar networks regimes under the most pronounced local desynchronization; it plays no role in the analyses reported in the main text.

All group-level statistics treat the subject as the unit of analysis. Quantities reported descriptively — component time courses, window means and source power — are averaged within a participant before the group mean is formed, so that every group mean and SEM is taken over the nine subjects. Where inference is performed the recordings are kept separate, but the permutation scheme respects the nesting: both sessions of a participant always receive the same sign, and all 2^9^ = 512 sign assignments are enumerated exhaustively. No test treats two sessions of one participant as independent observations, and the smallest two-sided *p* the design admits is 2*/*512 = 0.0039.

Since individual MRIs are not available for the dataset, the source reconstruction was performed using a template anatomy (fsaverage) and a three-layer boundary element model. The cortical source space was defined with icosahedral spacing (ico2) with 324 sources and median nearest-neighbour spacing 12 mm. A forward solution was computed using a realistic BEM geometry with conductivities (0.3, 0.006, 0.3) S/m for scalp, skull, and brain, respectively. Each source was modeled with three orthogonal dipoles using cortical patch statistics for orientation estimation. The forward operator was reduced to two tangential orientations per source via SVD of each source’s lead-field columns. To facilitate subspace-based connectivity analysis, the sensor-space forward operator was further projected onto a reduced spatial subspace via SVD of **G**_2*D*_**G***^T^*with a relative energy threshold of 10^−3^, yielding approximately 50 virtual sensors. This reduction retains the components of the forward model above a relative energy of 10^−3^; all subsequent projection and coupling estimation steps are performed in the reduced space.

Source-space power distributions were estimated by applying a minimum-norm inverse (*λ* = 10) to the band-specific sensor-level cross-spectral matrix, yielding per-source power as the leading singular value of the 2 × 2 source-level cross-spectrum. These power maps characterized the spatial distribution of induced activity and served as context-dependent weights for the CD-PSIICOS projector, focusing leakage suppression on sources with the strongest task-related power modulations. Functional connectivity was estimated using the CD-PSIICOS method described in Section 2.3. Coupling time series were baseline-corrected by subtracting the mean coupling in the pre-stimulus interval (−0.5 to 0.0 s). Source pairs separated by more than 70 mm are referred to as distal and those separated by less than 30 mm as local throughout. The networks analysed below were not defined window by window but obtained from a group decomposition of all 18 recordings, described in Section 2.4.

As a stress test of the findings, the analysis was also carried out under deliberately unfavourable conditions: the data were projected onto the orthogonal complement of the CSP components that maximally separate pre-from post-stimulus power, removing the sensor-space subspace that carries the movement-related local power modulation, and only the recordings in which desynchronization survived that projection were retained. Appendix E describes the procedure and the distal networks obtained under it.

### 2.3. Connectivity estimation

To estimate functional connectivity from both simulated and empirical MEG data, we employed the Context-Dependent Phase-Shift Invariant Imaging of Coherent Sources (CD-PSIICOS) framework (Kleeva and Ossadtchi, 2025; Ossadtchi et al., 2018), which allows source-space cross-spectral interactions to be recovered from MEG data while suppressing spatial leakage and power-related confounds. This approach is particularly suitable for the present study, as it enables explicit separation of local, power-dominated synchrony from distal, interaction-specific phase coupling. An open-source implementation is available at https://github.com/dkleeva/CD-PSIICOS.

Let **x**(*f, t*) ∈ C*^M^* denote the sensor-space signal at frequency *f* and time *t*, generated by source activities *J_i_*(*f, t*) ∈ C as **x**(*f, t*) = ∑^N^_i=1_**g***_i_J_i_*(*f, t*) + **n**(*f, t*). The source-level cross-spectral coefficients are

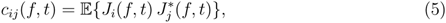

with *c_ii_* = E{|*J_i_*|^2^} ∈ R_≥0_ encoding source power and, for *i* ≠ *j*,

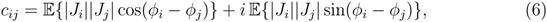

so that Re *c_ij_*captures instantaneous (zero- and near-zero-phase) interactions and Im *c_ij_* captures lagged interactions. The sensor-space cross-spectrum **C***_xx_*(*f, t*) = E{**x**(*f, t*)**x**^H^(*f, t*)} admits the vectorized representation

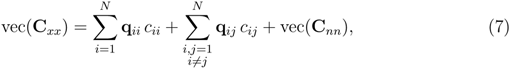

where **q***_ij_* = **g***_j_* ⊗ **g***_i_* are real-valued 2-topographies derived from the forward model. Since **q***_ij_* and *c_ii_* are real, this splits into

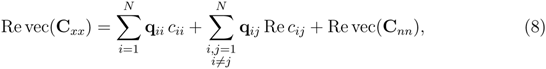

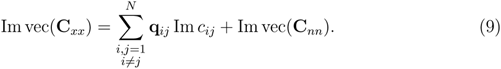

The auto-terms **q***_ii_c_ii_* span the spatial leakage subspace and contaminate only the real part; the imaginary part is leakage-free by construction — the basis of imaginary-coherency estimators, which however discard the instantaneous interactions encoded in Re *c_ij_*.

In the original PSIICOS approach (Ossadtchi et al., 2018; Altukhov et al., 2023) spatial leakage is suppressed by projecting vec(**C***_xx_*) onto the orthogonal complement of the subspace spanned by auto 2-topographies. Let

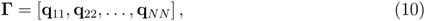

and let **U***_R_* denote the first *R* left singular vectors of **Γ**. The rank-*R* projection operator is defined as

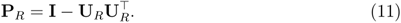

Applying **P***_R_* attenuates contributions dominated by source power and spatial leakage while preserving interaction-related components, including zero- and near-zero-phase coupling.

In CD-PSIICOS, the leakage subspace is adapted to the empirical cortical power distribution (Kleeva and Ossadtchi, 2025). Source power estimates *p_i_* obtained via a linear inverse operator are used to construct the weighted leakage matrix

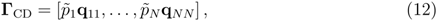

where *p*^~^*_i_* = *p*_i_/∥P∥. The corresponding projection operator

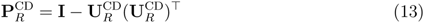

suppresses leakage originating from strongly active but uncoupled sources more efficiently, typically at lower projection ranks.

For a source pair (*i, j*), let **A***_i_* ∈ R*^M^*^×2^ contain the two tangential lead-field columns of source *i*, each normalized to unit norm. The projected source-level cross-spectral block is

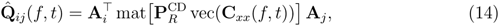

and the coupling score is its largest singular value, *c*^*_ij_*(*f, t*) = *σ*_max_(Re **Q**^^^ *_ij_* (*f, t*)), which aggregates over the two tangential orientations without an explicit orientation choice. This quantity is a cross-spectral coefficient expressed in units of source power: it scales with the product of the source amplitudes and with the consistency of their phase relation (Eq. 6), rather than being normalized by the power of the two sources as in the conventional coherence. We retain the unnormalized form because source-level power estimates are themselves leakage-contaminated, and because the projection removes the auto-term subspace in which source power resides, so that power normalization is not consistently defined in the projected space; normalized coherence between uncoupled sources moreover increases when their amplitude decreases. The estimate therefore reflects both amplitude and phase consistency; the two contributions are separated exactly by the factorization given in (Section 2.6). In the absence of coupling the estimate fluctuates around the noise floor; as a norm of the projected block it involves no division and is defined for every pair with *i* ≠ *j*.

The projection rank *R* controls how strongly power-dominated spatially local components are suppressed relative to interaction-specific ones; the trade-off this involves and the behaviour of the estimator at both extremes are treated in the original formulations of the method (Ossadtchi et al., 2018; Altukhov et al., 2023; Kleeva and Ossadtchi, 2025). Figure 2 locates the operating point in the present data. At *R* ≤ 10 none of the strongest pairs is distal: they connect neighbouring sources and sit on the regions of strongest band-limited power, the estimate still reading out the power topography rather than any interaction. Distal pairs enter at *R* = 20, occupy the ranking entirely by *R* = 50, and the composition does not change thereafter until *R* approaches the dimensionality of the whole auto-topography subspace, where interaction-specific structure is removed along with the power terms and the coupling estimates decline (Section 2.5). The rank used throughout, *R* = 300, falls well inside that plateau.

**Figure 2:**
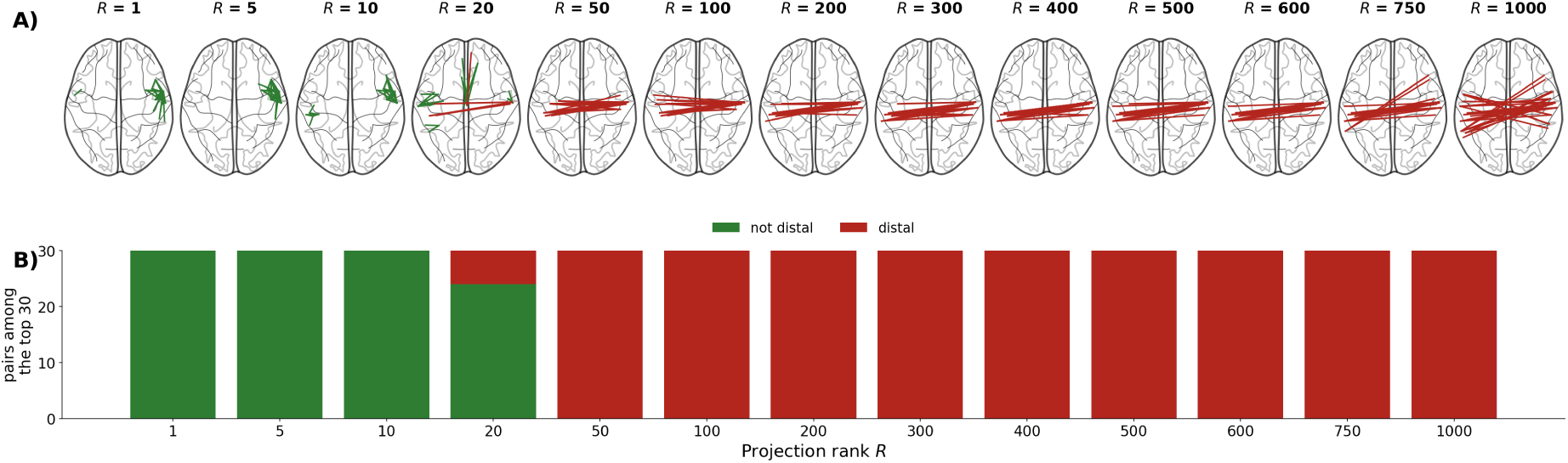
The selectivity of projection rank. (A) The thirty pairs with the strongest grand-average baseline-corrected coupling in the alpha 0.4–0.6 s window at each of the thirteen ranks of the sweep, red where the two sources are 70 mm apart or more and green otherwise. Each recording is divided by its own median absolute coupling at that rank before averaging, so that the ranking is not set by the few recordings with the largest amplitude. (B) How many of the thirty strongest pairs are distal at every rank.

### 2.4. Group decomposition

The networks analysed in the empirical part of this study were obtained from a decomposition of the coupling time courses of the whole sample. The input was the set of baseline-corrected coupling estimates at the fixed projection rank *R* = 300, one time course per source pair per recording, covering all 52 326 pairs of the 324-source space over the interval from −0.5 to 3.5 s. We chose a decomposition of this kind because the question asked here is not which networks are strongest on average, but which regimes of the relation between long-range and local coupling occur at all. A decomposition selects the pairs whose coupling covaries in time across the sample, so that each component is a candidate network whose reproducibility across participants can be tested explicitly, and networks belonging to different regimes are recovered separately rather than merged.

Decompositions of this kind have an established use in M/EEG connectivity research: temporal independent component analysis of envelope and time-resolved connectivity data has been used to recover task-and rest-related networks from MEG (Brookes et al., 2011; O’Neill et al., 2017), and state-space alternatives — hidden Markov models of fast transient networks (Baker et al., 2014; Vidaurre et al., 2017) and their continuous-mixture successors such as DyNeMo (Gohil et al., 2024) — as well as dynamic mode decomposition (Brunton et al., 2016) address the same problem of recovering group-level structure from heterogeneous recordings. We chose temporal ICA over the state-space alternatives because it makes no assumption that the data pass through a small number of discrete or mutually exclusive states.

For each component, we estimated the spatial profile of its network as the set of top 30 source pairs carrying the largest absolute mixing weight. Its window is the 0.2 s interval centred on the largest positive deflection of its group-mean time course, restricted to the post-stimulus interval. Its local counterparts are all local pairs (*<* 30 mm) whose two endpoints both belong to the sources used by that network. Neither the windows nor the pairs were selected by inspection, and the same rule was applied to every component in both bands.

Two requirements governed which components were described further: reproducibility under independent refitting in each half of a subject-wise split of the sample, and a coupling response inside the component’s own window reaching 0.8 of the pre-stimulus standard deviation. The stimulus-locking of each retained component’s peak was then tested against circular-shift surrogates, and its local load *E_L_* – the mixing weight it places on the short connections of its own sources, relative to all other short connections — was referenced to a null of random source sets. Appendix D describes all of these measures in full and reports their outcome for every component of both bands (Figs. 19 and 20, Tables 1 and 2).

**Table 1:** Alpha band: component selection and the surrogate test of the coupling peak. For every one of the twelve temporal-ICA components the table gives the two quantities on which the selection was based and, for the selected components, the test of the coupling peak. |*r*| is the median correlation between the component’s spatial map and its match when PCA and ICA are refitted independently in each half of a subject-wise split of the sample; *recovered* is the percentage of halves in which that match exceeded 0.7, with its Wilson 95% confidence interval. A component was considered reproducible when the lower bound of that interval exceeded 50%, the rate above which a component carried by a single participant cannot rise. *Distal response* is the mean coupling of the component’s 30 strongest pairs inside its own window, averaged over the nine subjects and expressed in units of each pair’s own prestimulus standard deviation; the second criterion required it to reach 0.8. Components meeting both criteria are shown in bold and were retained for further analysis. For these, the window is the 0.2 s interval centred on the largest positive deflection of the component’s group time course, and *p* is the fraction of 2000 circular-shift surrogates whose group mean in that window reaches the observed value; the surrogates preserve the amplitude, shape and autocorrelation of every recording and destroy only the alignment to the stimulus.

| Component | $\vert r\vert $ | Recovered | Distal response | Window (s) | $p$ |
| --- | --- | --- | --- | --- | --- |
| **IC1** | 0.71 | 61% [50, 71] | +1.90 | 0.31–0.51 | 0.0065 |
| IC2 | 0.71 | 51% [40, 62] | −0.79 | – | – |
| **IC3** | 0.91 | 85% [76, 91] | +1.20 | 2.16–2.36 | 0.0095 |
| **IC4** | 0.73 | 76% [66, 84] | +1.47 | 3.11–3.31 | 0.0010 |
| **IC5** | 0.74 | 61% [50, 71] | +0.91 | 2.36–2.56 | 0.0050 |
| IC6 | 0.86 | 56% [45, 67] | +1.17 | – | – |
| IC7 | 0.83 | 57% [47, 68] | +1.14 | – | – |
| IC8 | 0.77 | 75% [65, 83] | +0.12 | – | – |
| IC9 | 0.84 | 50% [39, 61] | +0.67 | – | – |
| IC10 | 0.76 | 50% [39, 61] | +0.22 | – | – |
| IC11 | 0.79 | 50% [39, 61] | +1.11 | – | – |
| IC12 | 0.81 | 50% [39, 61] | +0.90 | – | – |

**Table 2:** Beta band: component selection and the surrogate test of the coupling peak. Columns as in Table 1.

| Component | $\vert r\vert $ | Recovered | Distal response | Window (s) | $p$ |
| --- | --- | --- | --- | --- | --- |
| IC1 | 0.81 | 50% [39, 61] | +6.05 | — | — |
| IC2 | 0.81 | 51% [40, 62] | +6.95 | — | — |
| IC3 | 0.83 | 50% [39, 61] | +15.99 | — | — |
| IC4 | 0.65 | 50% [39, 61] | +17.38 | — | — |
| **IC5** | 0.93 | 76% [66, 84] | +6.57 | 1.16–1.36 | 0.0485 |
| **IC6** | 0.88 | 74% [63, 82] | +1.55 | 2.11–2.31 | 0.0115 |
| IC7 | 0.78 | 64% [53, 73] | +0.51 | — | — |
| IC8 | 0.57 | 25% [17, 35] | +0.41 | — | — |
| IC9 | 0.56 | 9% [4, 17] | +0.45 | — | — |
| IC10 | 0.81 | 91% [83, 96] | +0.73 | — | — |
| IC11 | 0.83 | 76% [66, 84] | +0.63 | — | — |
| IC12 | 0.93 | 95% [88, 98] | +0.54 | — | — |

### 2.5. Subpopulation analysis of local coupling trends

To assess how local coupling among distally connected nodes evolves with progressive leakage suppression, the CD-PSIICOS projection was applied at 13 ranks (*R* = 1, 5, 10, 20, 50, 100, 200, 300, 400, 500, 600, 750, 1000). For each rank and each local pair, the window-mean of the coupling time course was extracted and normalized by the maximum absolute value of the group mean across all local pairs at that rank, yielding one value per pair per rank. A linear slope fitted across ranks classifies the pair as *leakage-resistant* (positive slope, coupling preserved or enhanced after power-leakage removal) or *leakage-driven* (negative slope, apparent coupling attributable primarily to spatial leakage). In the beta band the contribution of the transient components was removed at every rank before this step, as described in Section 2.4.

Because a linear slope is a coarse summary of a non-monotonic curve, we do not attach a significance test to the trend of either subpopulation separately and report only the contrast between them, together with the level at which the local pairs settle. The level of a pair is its window-mean coupling averaged over the high ranks (*R* ≥ 200), that is, the range in which the projector has removed the bulk of the power-dominated subspace and the local estimates no longer change systematically with rank, expressed in units of that pair’s own pre-stimulus standard deviation at each rank. It is summarised by the median over all local pairs of a network, by the median within each subpopulation, and by the number of pairs lying above the pre-stimulus level.

The contrast was computed per recording as the mean slope of the leakage-resistant pairs minus the mean slope of the leakage-driven pairs, giving 18 values. All 18 recordings were retained, and the non-independence of a participant’s two sessions was handled by the test rather than by averaging: sign flips were applied at the level of the nine subjects, so that both sessions always receive the same sign (2^9^ = 512 permutations, two-sided, smallest attainable *p* of 0.0039). Effect size is the median difference with a 95% confidence interval bootstrapped over subjects. Because the subpopulations are defined by the sign of the grand-average trend, their separation is positive by construction; the test asks only whether individual recordings reproduce it consistently, and provides no inferential evidence for the networks themselves.

### 2.6. Phase alignment analysis

The coupling reported throughout is the squared largest singular value of Re *B*^-^*_ij_* (*t*), where *B*^-^*_ij_* (*t*) is the 2 × 2 cross-spectral block of the pair (*i, j*) after the CD-PSIICOS projection. This is a magnitude, and it can grow either because the single-trial cross-terms grow or because their phases agree better across trials. The two contributions separate exactly, because the projector is a linear operator on the vectorised cross-spectrum and therefore commutes with the average over trials: *B*^-^*_ij_* (*t*) = ⟨*B_ij,k_*(*t*)⟩*_k_*, where *B_ij,k_*(*t*) is obtained by applying the same rank-300 projector to the cross-spectrum of the single trial *k*.

Let *u*(*t*) and *v*(*t*) be the leading left and right singular vectors of Re *B*^-^*_ij_* (*t*), so that the reported coupling is (*u*^⊤^Re *B*^-^*_ij_ v*)^2^, and let *s*_k_ (*t*) = *u*(*t*)^⊤^*B_ij,k_*(*t*) *v*(*t*) be the corresponding single-trial scalar, with modulus |*s_k_*| and argument *φ_k_*. Since *s_k_* is linear in *B_ij,k_*, the coupling factorises as

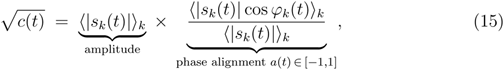

which we verified numerically against the coupling produced by the main pipeline. The alignment *a*(*t*) is an amplitude-weighted circular mean of the single-trial phases of the leakage-free cross-term: it approaches 1 when every trial contributes at the same phase and 0 when the trial phases are uniform. It is estimated across trials separately at each time point, so a window enters only as an average over per-time-point estimates, and the length of the window is not a source of bias.

For each component both factors were averaged over its top pairs and over the time points of the interval of interest, giving one value per recording. The interval of interest was a 400 ms window centred on the window used elsewhere in the analysis, and the reference was the prestimulus interval (−0.4 to 0 s). All 18 recordings were kept, and the dependence between the two sessions of a participant was handled by the test rather than by averaging: the permutation distribution was generated by exhaustive sign flips applied at the level of the nine subjects. The singular vectors *u*(*t*), *v*(*t*) are read off the data at each time point, which raises the absolute level of *a*(*t*), but identically in the prestimulus interval and in the window, so it cannot create a difference between them.

## 3. Results

### 3.1. Simulations

We use the controlled realistic simulations to verify that the regime targeted by the empirical analysis, long-range phase coupling that persists while within-region synchrony is destroyed, is recoverable from sensor-level MEG given a leakage-aware connectivity pipeline, when such a regime is present in the generating model. The simulation framework is designed to satisfy two properties simultaneously: (i) vertex amplitudes are held strictly constant, so any change in MEG-observable signal traces back to phase coherence alone; and (ii) the paired-jitter mechanism guarantees that distal phase relationships between corresponding subpopulations across hemispheres remain intact regardless of within-region phase dispersion.

Across the within-region jitter sweep the mean sensor power fell by roughly an order of magnitude and saturated at the plateau set by the surviving paired cross-region component, with the knee at *σ*_local_ ≈ 0.5*π*, and the focal sensor topography of the two ROIs collapsed over the same range. No parameter of the source signal changed: every cortical vertex emits a sinusoid of the same fixed amplitude *A* throughout, so the whole effect is a cancellation between subpopulations at the macroscopic level. A representative realisation is shown in Appendix A (Fig. 13).

To establish the operating regime of the projection rank we inspected the distance profile of pair-wise coupling across ten ranks from *R* = 0 to *R* = 700 at *σ*_local_ = 0, (Appendix A, Fig. 14). Without projection it peaks at the nearest-neighbour bin, the signature of spatial leakage; from *R* = 1 onwards the maximum sits instead at the ground-truth inter-ROI distance of 10.8 cm, so the surviving cross-spectrum is interaction-specific and the choice of rank is forgiving; we use *R* = 100 in the remaining simulations. Crucially, the bias of CD-PSIICOS toward distal interactions is the reason that all subsequent comparisons in this paper with respect to simulations are made *within* a connection class rather than between local and distal absolute magnitudes. The relevant test of the hypothesis is whether the distal coupling that CD-PSIICOS isolates remains stable under within-region jitter, not whether it exceeds the projection-suppressed local component, which by construction it always will.

To test how the assumed microarchitecture of the populations affects the recovery of long-range coupling, we swept the subpopulation count *K* ∈ {2, 5, 10, 20, 30} at fixed Δ = *π/*20 and tracked the source-level distal coherent sum *S*^dist^ across the jitter range (Appendix A, Fig. 15). Each curve decays smoothly and saturates by *σ*_local_ ≈ *π* on the theoretical 1*/K* plateau, from 0.50 at *K* = 2 to 0.033 at *K* = 30, so the fraction of distal coupling that survives within-region jitter is an inverse function of how finely the cortex is subdivided into independent oscillatory units. That decay does not make distal coupling undetectable relative to local coupling, however: the local coherent sum collapses entirely once within-region phase coherence is destroyed, while the distal sum only falls to its plateau, so the distal-to-local ratio grows with *σ*_local_ rather than shrinking, and even the 3.3% remaining at *K* = 30 still exceeds the within-ROI residual. To compare the leakage-aware CD-PSIICOS pipeline against the standard imaginary-coherence baseline, we ran both estimators on identical simulated MEG data for three values of the subpopulation count, *K* ∈ {5, 10, 30} and the full within-region jitter sweep *σ*_local_*/π* ∈ [0, 2]. For each *K* we report six observables side by side (Figure 3). At every *σ*, the distal class is the larger of the two coherent sums for both estimators.

**Figure 3:**
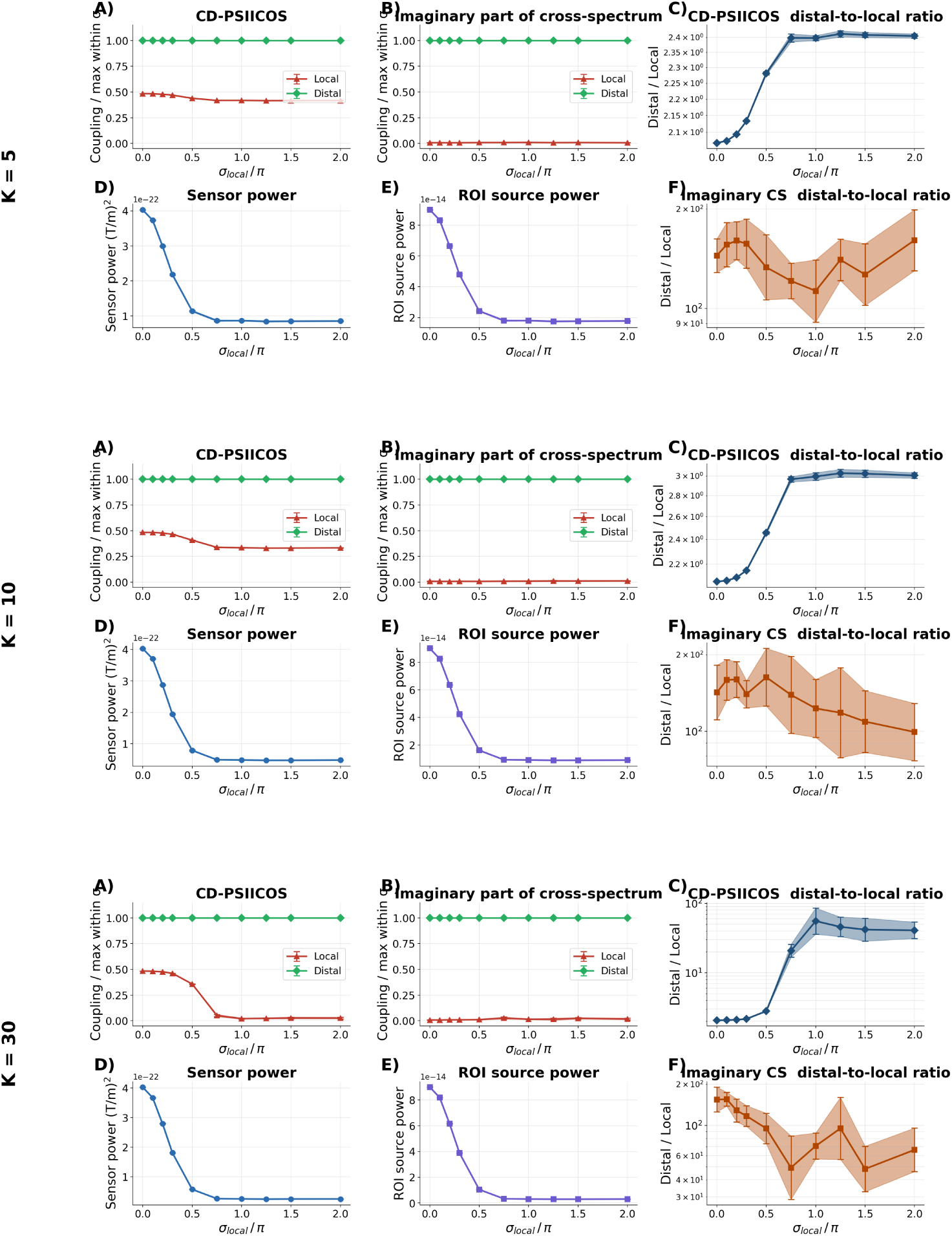
CD-PSIICOS vs imaginary-coherence baseline across subpopulation counts *K* ∈ {5, 10, 30}. Each row is one *K* (label on the left); within each row the six panels (A–F, repeated per row) are: (A) CD-PSIICOS coupling — within-*σ* normalized local (red) and distal (green) traces; (B) imaginary part of the cross-spectrum (no projection) — same normalisation; (C) CD-PSIICOS distal-to-local coupling ratio (log *y*); (D) mean sensor power (planar gradiometers); (E) mean ROI source power; (F) imaginary-CS distal-to-local ratio (log *y*). All curves show mean ± 95% CI across *N*_MC_ Monte-Carlo trials with *σ*_local_*/π* ∈ [0, 2], *f*_0_ = 10 Hz, *T*_sim_ = 600 s, Δ = *π/*20.

The informative quantity is therefore the local trace (red) — the relative magnitude of within-ROI coupling expressed as a fraction of the distal-or-local maximum. With CD-PSIICOS (Panel A), local sits between 0.4 and 0.6 at *σ* = 0 and decays with *σ*_local_; the rate of decay is set by the subpopulation count: at *K* = 5 it drops only to ≈ 0.4 (a substantial fraction of within-ROI pairs share a subpopulation and remain coherent), at *K* = 10 to ≈ 0.3, and at *K* = 30 — where each cortical vertex is its own subpopulation and no within-ROI pairs remain coherent — local collapses to *<* 0.05. With the imaginary cross-spectrum baseline (Panel B), local stays near zero across all *σ* and all *K*: imaginary coherence cannot resolve the zero-phase within-ROI coupling that exists at *σ* = 0, because it discards the real-valued cross-spectral component by construction.

At the same time, both power observables decrease monotonically with *σ*_local_ (with overlapping CIs across *K* for sensor power, and clearly *K*-dependent ROI power that drops by a factor of *K* to its 1*/K* plateau by *σ*_local_ ≈ 0.75*π* in every condition, exactly as the generative model predicts). This confirms that the macroscopic power signal that conventional sensor-level analyses depend on collapses by an order of magnitude across the jitter sweep.

Together, the results show that CD-PSIICOS is not biased toward distal coupling in a way that obscures local desynchronization; instead it is the unique estimator (among those compared) whose distal-to-local ratio is sensitive to within-region jitter, and therefore the appropriate choice for testing whether long-range phase coupling is preserved when local synchrony is destroyed.

This comparison depends on the choice of inter-regional lag. The simulations above use a single inter-regional lag Δ = *π/*20, and ImCS is by construction blind to zerolag interactions, so it suppresses every within-ROI pair of our generative model, whose mean phase difference is zero, while cross-ROI pairs, differing by Δ ≠ 0, survive. ImCS is therefore structurally biased *toward* the distal class in this benchmark, which is what makes it appear competitive and what fails the moment the distal lag itself approaches zero: at *δ* = 0 its PR-AUC falls to ≈ 0.2 against ≈ 0.61 for CD-PSIICOS, which holds that plateau until its own mirror failure point at *δ* = *π/*2. Appendix B reports the full phase-lag and jitter comparisons and the per-pair detection curves at the two contrasting lags.

### 3.2. Real data analysis

#### 3.2.1. Event-related desynchronization and power modulations

To explore the presence of the dissociation between local desynchronization and long-range phase coupling we used the data from empirical MEG recordings acquired during voluntary movement. Using a center-out reaching paradigm, we quantified movement-related power modulations and source-space phase coupling to assess how distal interactions evolve under conditions of pronounced local desynchronization.

Time–frequency analysis of planar gradiometer signals revealed robust movement-related desynchronization in both the alpha/mu (8–13 Hz) and beta (15–25 Hz) frequency bands, which were therefore selected for all subsequent analyses. In both bands power decreased shortly after movement onset, reached a minimum approximately 0.8–1.2 s later and then rebounded gradually. The suppression was not restricted to sensorimotor sensors: in the alpha/mu band it extended toward posterior sensors, indicating additional involvement of occipital regions, with a similar but weaker posterior contribution in the beta band. The grand-average time–frequency representation, the band-limited power time courses over the left and right parietal sensor groups, and the topographic series on which these observations rest are presented in Appendix C (Fig. 18).

Source-space power distributions were next examined. Figure 4 summarizes the grand-average source-level power modulations in the alpha and beta bands, reconstructed using a minimum-norm inverse solution and averaged across the selected sessions.

**Figure 4:**
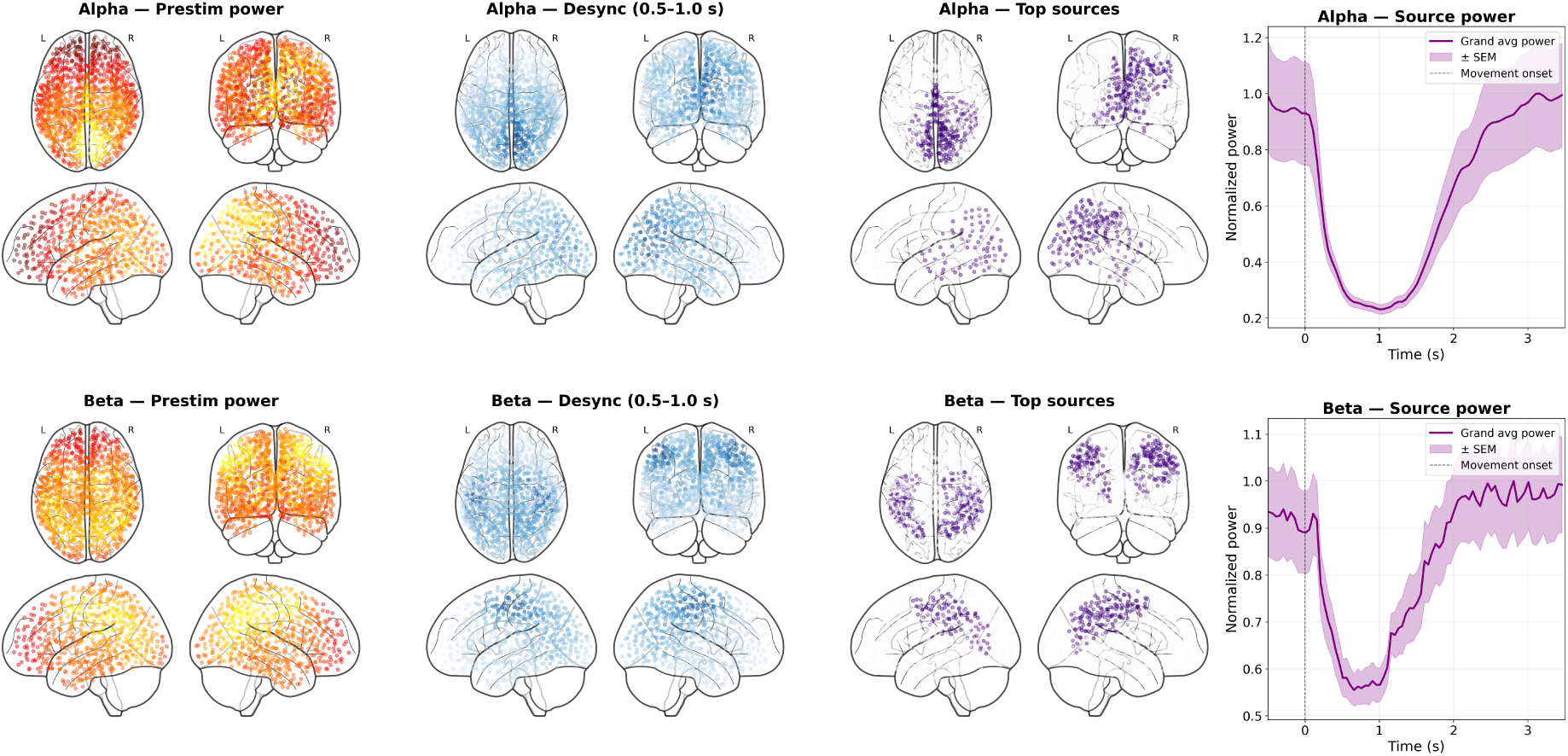
Grand-average source-space power modulations. (A) Alpha band (8–13 Hz). (B) Beta band (15–25 Hz). For each band, from left to right: prestimulus source power distribution; desynchronization map computed as relative power change in the 0.5–1.0 s post-movement window; cortical sources exhibiting the most pronounced desynchronization (purple dots); grand-average source power time course (±SEM across sessions and subjects, vertical dashed line marks movement onset).

In the alpha band (Figure 4A), the spatial distribution of source power was dominated by a focal posterior (occipital) cluster. The corresponding grand-average source power time series exhibited a pronounced decrease following movement onset, reaching a minimum during the movement execution period and gradually recovering thereafter.

In the beta band (Figure 4B), the source-level power distribution showed a broader spatial pattern. The elevated power modulation extended over more diffuse regions involving sensorimotor and parietal regions. The beta-band source power time course showed a marked reduction after movement onset, followed by a gradual rebound. Notably, the presence of a posterior alpha-dominant pattern together with a more spatially extended beta pattern is consistent with the sensor-level topographies observed in the time–frequency analysis (Figure 18).

#### 3.2.2. Distal networks in alpha band

Having established the presence of movement-related desynchronization in both frequency bands and characterized the source-space power distributions, we next examined distal phase coupling and its relationship to local synchronization at the fixed projection rank (*R* = 300). Time windows and the networks active within them were obtained from a group decomposition of the full sample. A temporal ICA of the concatenated coupling time courses of all 18 recordings yields *K* = 12 components per band, each consisting of a weight vector over all 52 326 source pairs and a single time course common to the group. For each component we define its network as the 30 pairs carrying the largest absolute weight, its window as the 0.2 s interval centred on the largest positive excursion of its group time course, and its local counterparts as the local pairs whose two endpoints both belong to that network’s nodes. The same procedure is applied to both frequency bands.

Four of the twelve alpha components met both criteria — IC1, IC3, IC4 and IC5 — and are described below. All twelve, with their networks, their group time courses and their local load, are shown in Appendix D (Fig. 19), which also gives the two selection quantities and the surrogate test of the coupling peak for every component (Table 1).

The first alpha component, IC1, peaks at *t* = 0.41 s. on an entirely interhemispheric network linking right supramarginal cortex with left postcentral and supramarginal cortex (Fig. 5A), where coupling rises by +1.90 prestimulus SD inside its window (Table 1). Over the same interval the short connections anchored to those sources move the other way, falling by −0.53 (Fig. 5B), while the power of the sources themselves falls by −3.60 (*p* = 0.031). Both hold in seven of the nine subjects, as does the distal–local separation of +2.43 (*p* = 0.020). The long-range interaction thus strengthens at the moment when the regions carrying it are most strongly desynchronized and their own short connections least coupled. IC1 is also among the alpha components that load least on short connections (*E_L_*= 0.56, below its null; Appendix D, Fig. 19B).

**Figure 5:**
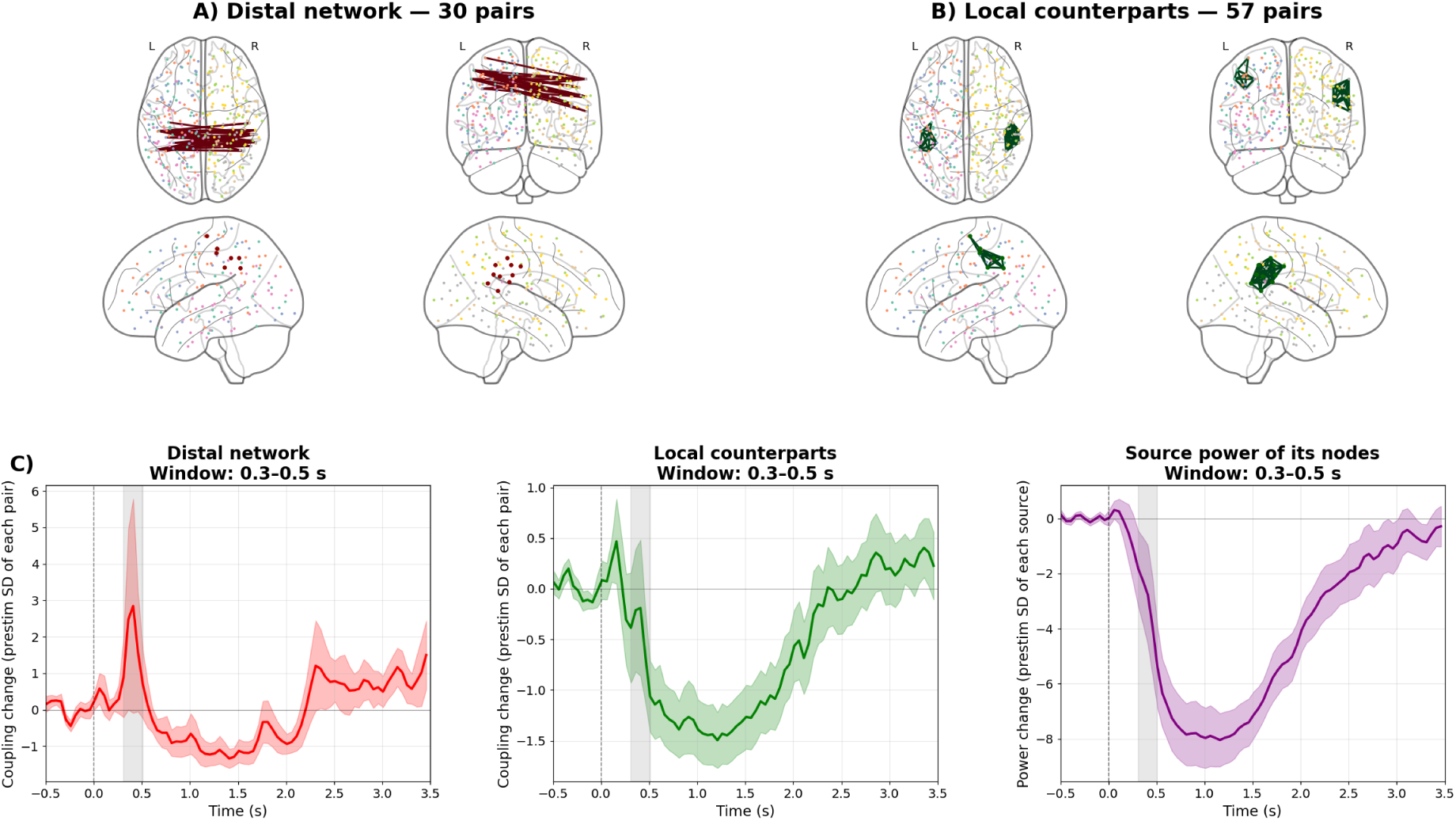
Alpha component IC1. (A) The network, in four views. (B) The local pairs whose endpoints are both its sources. (C) Coupling of the distal network, of the local pairs, and power of the sources, as the mean over nine subjects ± SEM, each in units of its own prestimulus standard deviation. Grey band: the component’s window, 0.31–0.51 s; dashed line: stimulus onset.

To ask whether the local pairs of this network retain any phase structure of their own, the CD-PSIICOS projection was recomputed at thirteen ranks, from *R* = 1 to *R* = 1000, and each local pair was classified by the sign of its grand-average trend across ranks. Ten pairs showed a positive trend and were classified as leakage-resistant, and forty-seven pairs showed a negative trend and were classified as leakage-driven (Fig. 6D). The two subpopulations separate in the expected direction: the difference between their mean per-recording trends is positive in all eighteen recordings, with a median difference of +0.047 (95% CI [+0.040, +0.064], bootstrapped over subjects; two-sided sign-flip test with flips applied at the level of the nine subjects, *p* = 0.004).

**Figure 6:**
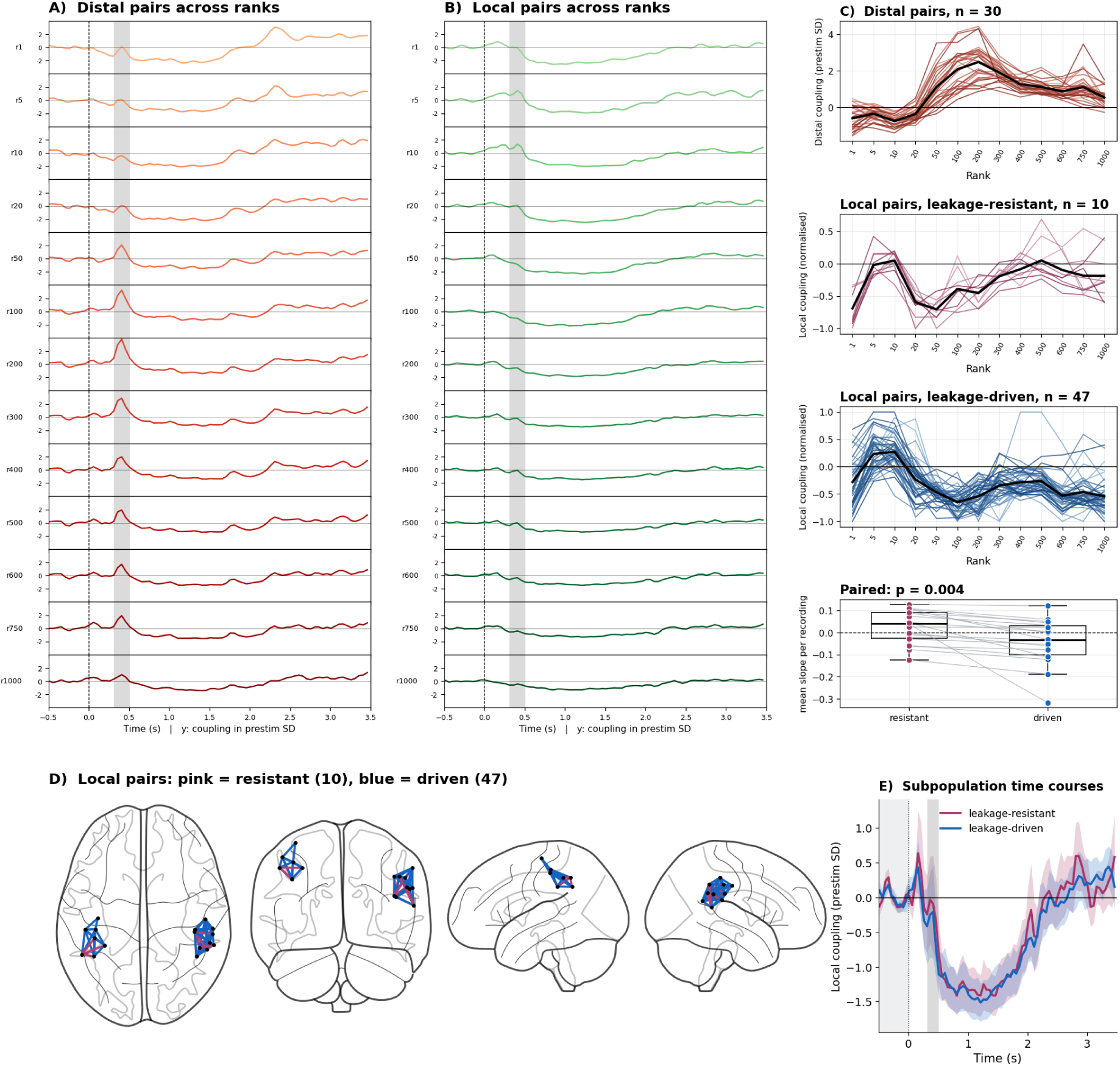
Rank dependence of the local pairs of alpha component IC1. (A, B) Coupling of the distal network and of the local pairs across the thirteen ranks, each in units of its own prestimulus standard deviation at that rank; grey band: the component’s window. (C) Window-mean coupling against rank for the distal pairs (top) and for the leakage-resistant and leakage-driven local pairs (middle, bottom); the local panels are normalised within each rank by the largest absolute group mean over all local pairs, which is what defines the classification. Box plot: mean trend per recording in the two subpopulations, with the two-sided sign-flip *p*. (D) The local pairs on the cortex, by classification. (E) The two subpopulations at *R* = 300, mean over nine subjects ± SEM.

The level at which the two subpopulations settle is even more informative than their trends. Averaged over its pairs, the leakage-resistant subpopulation stays below the prestimulus level at every rank at which the projector is active (R ≥ 50), with a single excursion above it, to +0.05, at one rank. Pair by pair over the high ranks (R ≥ 200) the picture is the same: only 4 of the network’s 57 local pairs settle above their own prestimulus level, against a median of −0.65 over all of them. Progressive suppression of power-related contributions therefore reduces the depth of the local decoupling in this subpopulation but does not lift it to the prestimulus level, so the distal interaction of IC1 is not accompanied by a locally synchronized subpopulation that could support it. The coexistence of a stimulus-locked distal coupling increase with a predominantly leakage-driven local subpopulation suggests that the alpha-band interhemispheric interaction in this time window does not require local oscillatory synchrony within the participating regions.

The preceding analysis of IC1 illustrates one regime in detail: a long-range interaction that strengthens while the short connections of the same sources are suppressed. Because the four alpha components differ in exactly this respect, we summarise all four together in Fig. 7. Each row shows the component’s network and its local pairs, the coupling of both and the power of the participating sources, how the two evolve as the projection rank is increased, and where the local pairs finally settle relative to the prestimulus level.

**Figure 7:**
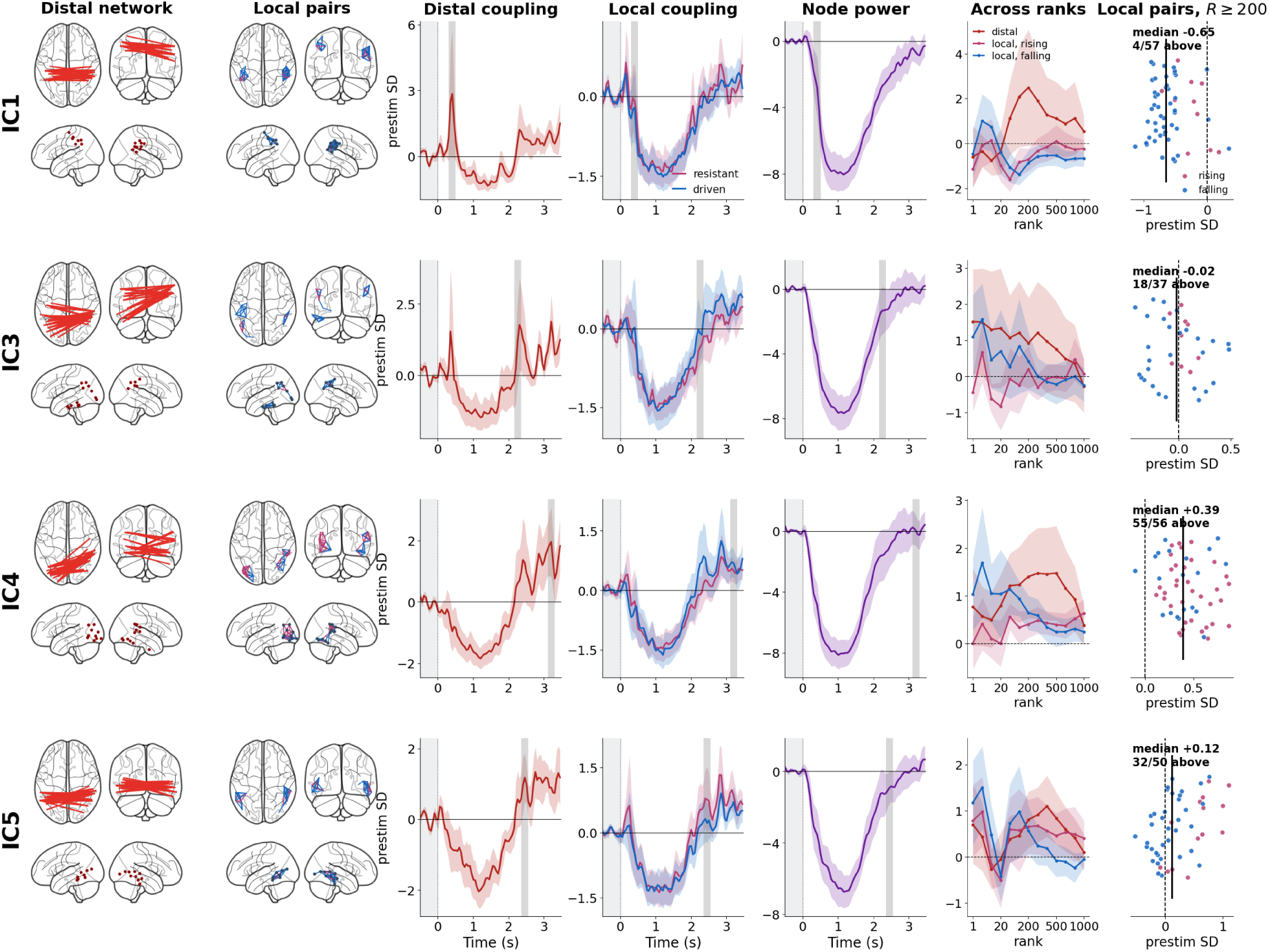
The four alpha networks, one per row. From left to right: the network in four views; its local pairs, coloured by whether their coupling rises (pink) or falls (blue) with projection rank; the coupling of the distal network; of the two local subpopulations; the power of the sources; the window-mean of all three against rank; and the level each local pair settles at for *R* ≥ 200, with the prestimulus level dashed, the median over pairs in black, and that median and the number of pairs above prestimulus in the title. Units and averaging as in Fig. 5.

IC3 connects right supramarginal cortex with left inferior parietal and inferior temporal regions and peaks at *t* = 2.26 s, by which time the power of its sources has recovered most of its drop (−1.34 prestimulus SD). Only 9 of its 37 local pairs are leakage-resistant, and they stay flat around the prestimulus level across the whole range of ranks (median +0.03 at high ranks), while the remaining 28 start at +1.09 and decay through zero, as an estimate made of leakage should. The cloud therefore straddles baseline (median −0.02, 18 of 37 above it): this network has neither the suppressed local coupling of IC1 nor a locally coherent subpopulation of its own, and what little survives locally lies in the parietal patches rather than in the ventral cluster where over half of its long-range pairs terminate (16 of 30). Its long-range estimate is stable rather than growing, +1.51 at *R* = 1 against +1.20 at *R* = 300, so IC3 is the only one of the four that does not strengthen as leakage is removed, and we treat it as the least conclusive of them.

IC4 links left lateral occipital cortex with right inferior temporal, fusiform and supramarginal regions across 22 sources and peaks last of the four, at *t* = 3.21 s, rising by +1.47 prestimulus SD inside its window (Table 1). By then the power of its sources has returned to the prestimulus level (−0.08 SD, a 99% recovery from the minimum of −8.15 at 1.01 s), so unlike the other three this is a network which results from the cortical activity that is no longer desynchronized.

However, its local pairs are the point of interest. 36 of 56 pairs are leakage-resistant, and here the classification separates two populations that both sit above baseline rather than one above and one below: their median levels at high ranks are +0.41 and +0.37 prestimulus SD, with 55 of the 56 pairs above the prestimulus level. The resistant subpopulation grows with the projection rank, from +0.01 at *R* = 1 to +0.64 at *R* = 1000, while the remainder decays from +1.04 towards +0.24 without crossing baseline. Removing the power-dominated subspace thus leaves the short-range coupling of these sources intact and, for most pairs, more prominent, which points to genuine macroscopic local phase alignment rather than the shadow of a power change.

Crucially, the long-range estimate behaves in the same way rather than in the opposite one. It strengthens monotonically over the working range of ranks, from +0.79 at *R* = 20 to +1.48 at *R* = 500, nearly doubling as leakage is suppressed, which is the signature of an interaction carried by phase rather than by power. Both ends of the spatial scale are thus leakage-resistant in this network, and they rise together. The most economical reading is that the long-range interaction of IC4 is accompanied, and plausibly supported, by locally synchronized activity in the participating regions. Since the local and distal coupling in IC4 grows rather shrinks as the power-dominated subspace is removed, source power alone is not what produces this picture.

IC5 spans bilateral middle temporal cortex and the banks of the superior temporal sulcus and peaks at *t* = 2.46 s, rising by +0.91 prestimulus SD inside its window (Table 1). Unlike IC4 it peaks while its sources are still desynchronized: their power has recovered 86% of its drop but remains 0.93 SD below baseline.

Its local pairs split more sharply than in any other alpha network. Thirty-six of the fifty start at +1.18 prestimulus SD and are driven to zero by *R* = 500, while the remaining fourteen hold between +0.4 and +0.7 across the entire range of ranks, with a median at high ranks of +0.62 against +0.03 for the others. Most of what is measured on the local connections of these sources is therefore leakage, removed together with the power subspace, while a minority retains phase alignment the projection does not affect. The long-range estimate strengthens over the same range, from −0.05 at *R* = 20 to +1.10 at *R* = 400, so it is leakage-resistant in its own right.

IC5 thus realises the intermediate configuration: a small locally coherent subpopulation survives inside cortex whose power is still well below baseline, and the long-range interaction strengthens alongside it.

Ordered by the time at which they peak, the four alpha networks trace a single progression: the more the power of the participating sources has recovered, the more of the local structure among those sources survives the projection. A long-range response is present throughout. The band therefore contains both extremes: at one end is coupling at distance with almost no macroscopic local synchronization to to support it; at the other, coupling at distance supported by local phase alignment accompanying the end of macroscopic ERD. The remaining two networks lie between them.

#### 3.2.3. Distal networks in beta band

The same decomposition of the beta band returns twelve components, of which two — IC5 and IC6 — meet both criteria; the other ten, and the quantities on which the selection rests, are given in Appendix D (Fig. 20, Table 2).

The first four components require the separate comment, because they capture sharp transients: each is confined to a fraction of a second, and is strongly loaded on the short connections of its own sources (*E_L_* between 1.59 and 2.77). None of them passes the reproducibility criterion. They matter nonetheless, because their spatial maps overlap with those of the components we do describe, so the pairs of any one network also carry a share of these events and their raw coupling shows a spurious excursion at the latency of the transients rather than of the network itself. For the beta networks below we therefore show the coupling of their pairs both in the raw data and after the contribution of IC1 to IC4 has been removed.

IC6 is a bilateral sensorimotor network, linking left postcentral with right supramarginal and right postcentral cortex across 17 sources, and peaks at *t* = 2.21 s, rising by +1.55 prestimulus SD inside its window (Table 2).

The raw time courses contain one feature that does not belong to this network. Between 0.8 and 1.05 s the local pairs of these sources rise to +1.42 prestimulus SD, higher than any local value IC6 reaches in its own window, and the distal pairs to +0.80 (Fig. 8C). The peak falls at the latency of the transient components IC1 to IC4, more than a second before that of IC6, and disappears once their contribution is removed (−0.26 and −0.36 over the same interval; Fig. 8D). The window of IC6 itself is unaffected either way: the distal estimate changes from +1.55 to +1.52 and the local one from +0.20 to +0.18. Nothing that follows therefore depends on the subtraction; it removes an event belonging to two other recordings from an interval in which this network has no peak of its own.

**Figure 8:**
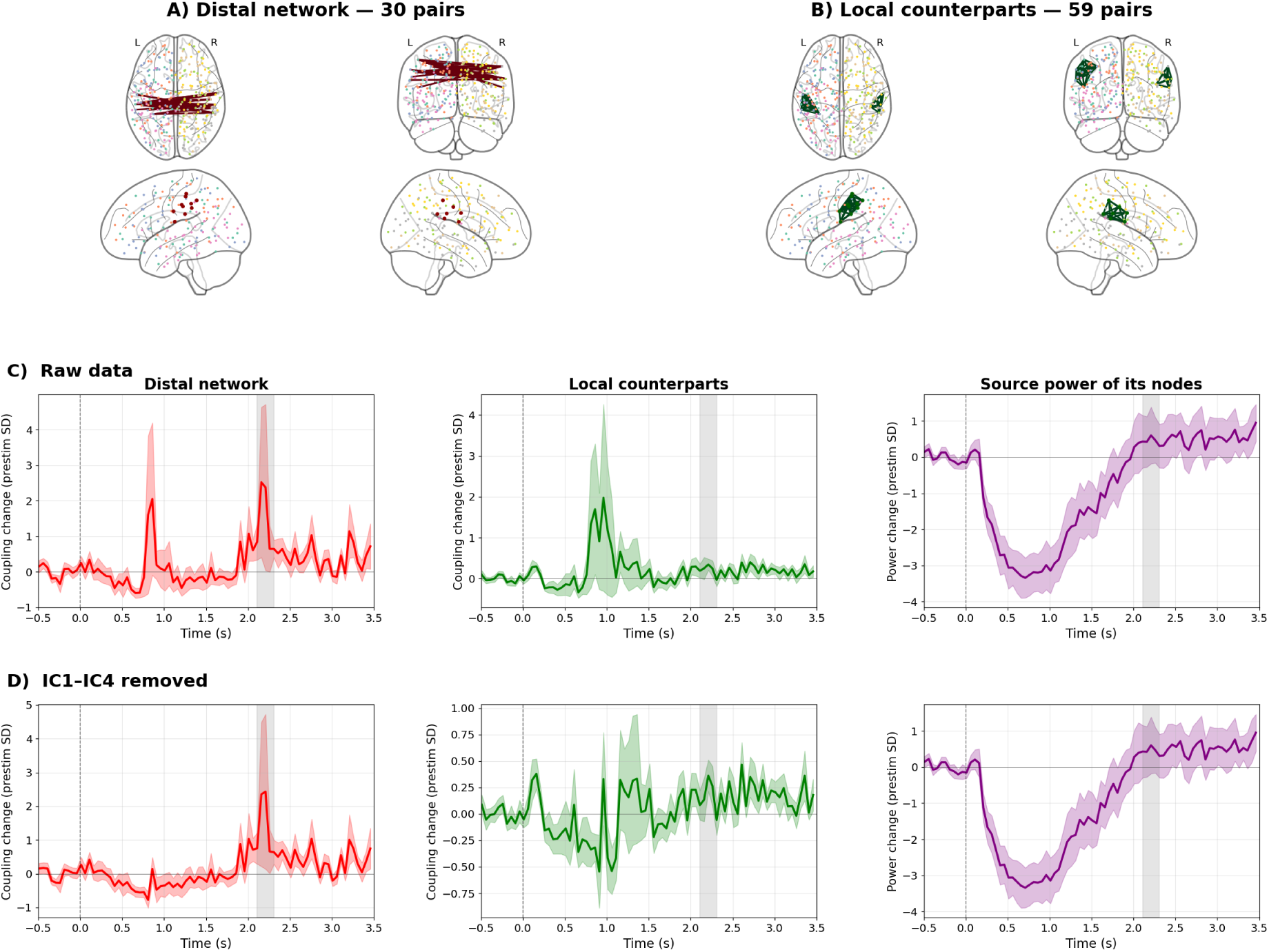
Beta component IC6. Layout and units as in Fig. 5, with (C) the raw data and (D) the same quantities after the contribution of the transient components IC1–IC4 has been removed; the deflection near *t* ≈ 0.9 s in (C) is that contribution. Window 2.11–2.31 s.

**Figure 9:**
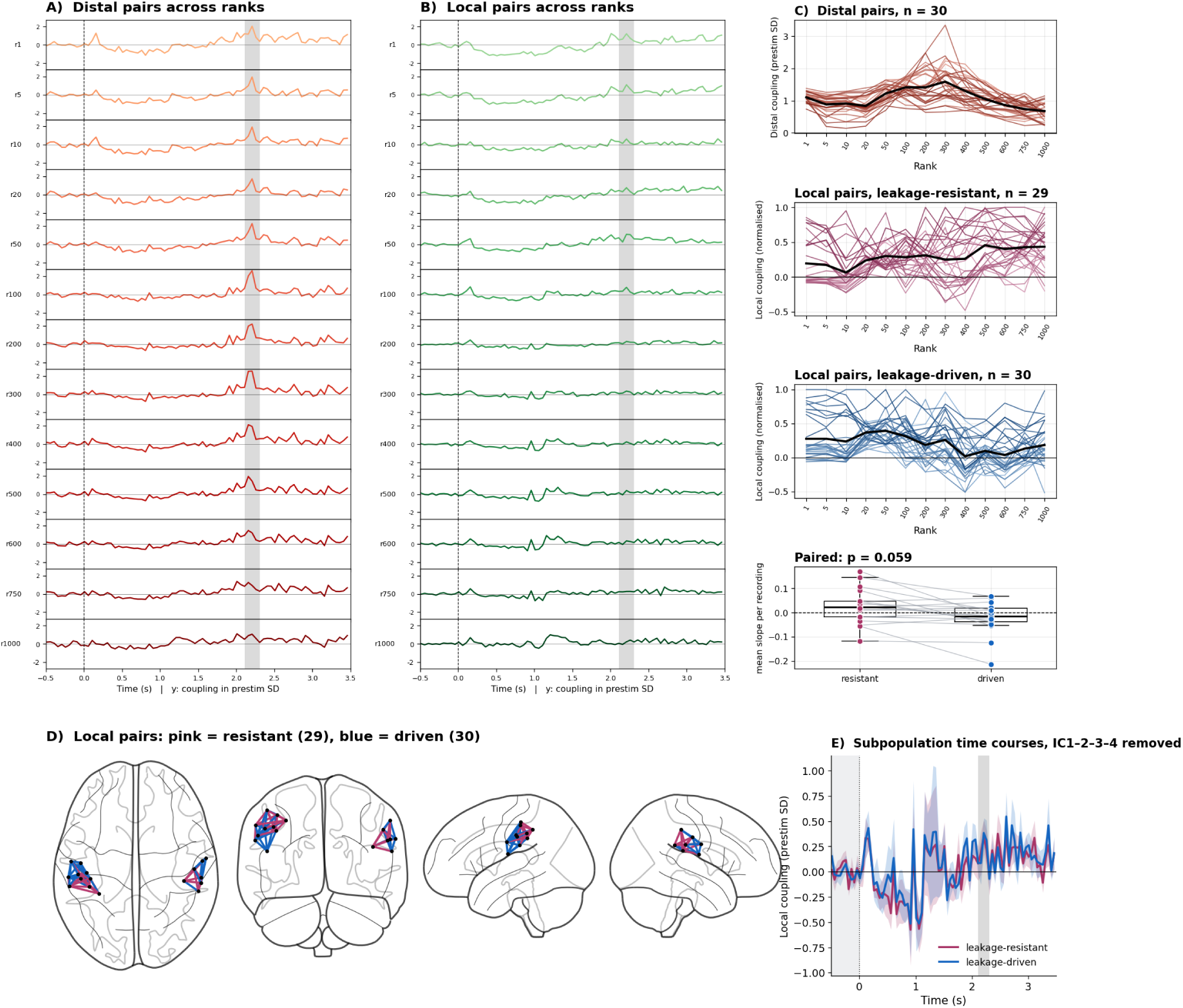
Rank dependence of the local pairs of beta component IC6. Panels and units as in Fig. 6, with the contribution of the transient components IC1–IC4 removed at every rank; the raw time courses are in Fig. 8C.

**Figure 10:**
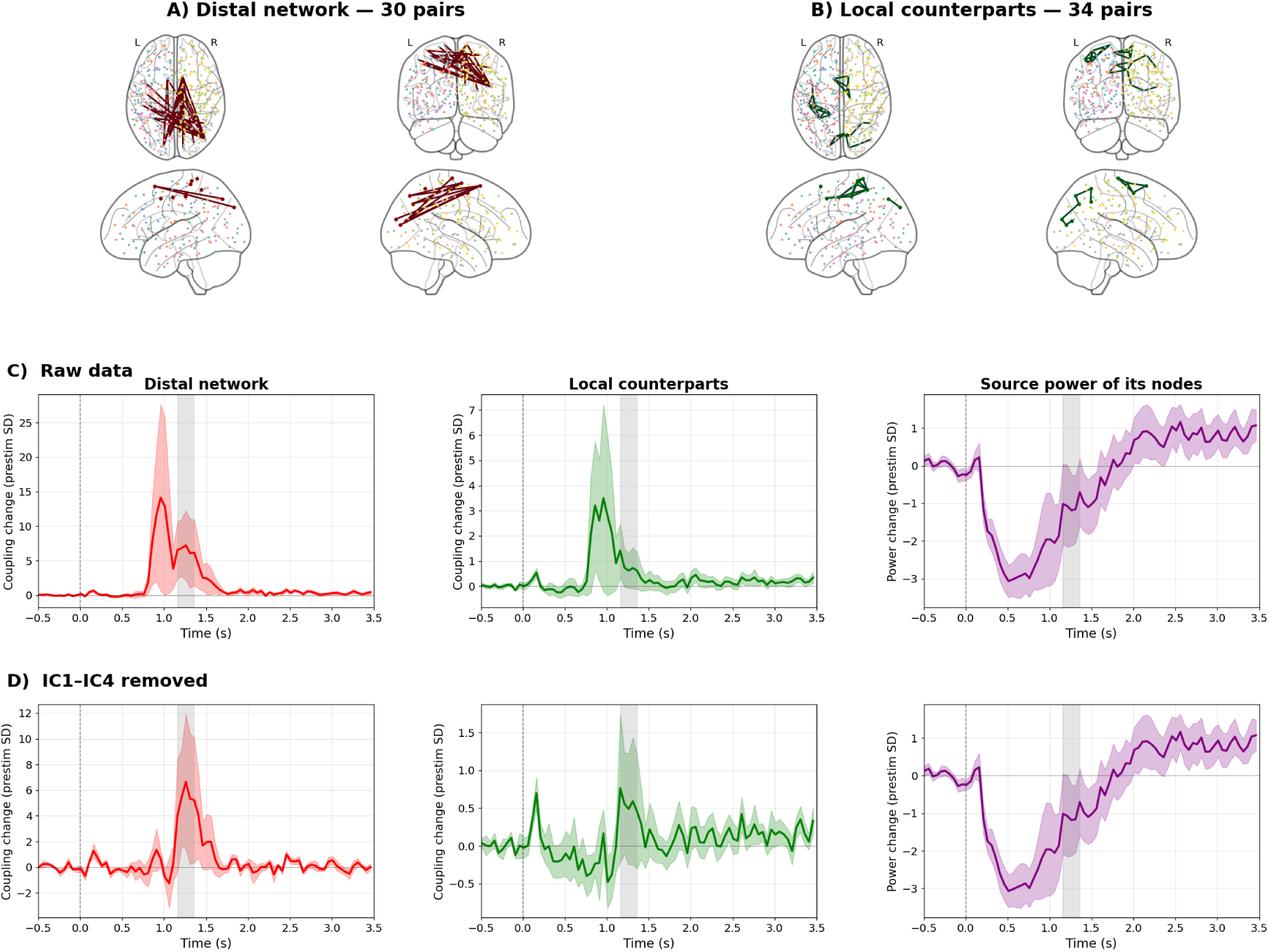
Beta component IC5. As in Fig. 8: (C) raw, with the first peak near *t* ≈ 0.9 s the contribution of IC1–IC4 and the peak of IC5 itself at 1.26 s; (D) after that contribution is removed. Window 1.16–1.36 s.

The timing of IC6 places it well into the beta rebound, and its peak must not be conflated with the recovery of power in the same sources. Power returns gradually and monotonically over the whole post-movement interval, from −3.34 prestimulus SD at *t* = 0.71 s to +0.28 at 2.01 s, drifting slowly upward thereafter without an event of its own. The distal coupling behaves differently: it stays at or below the prestimulus level throughout that recovery, rises abruptly to +2.43 prestimulus SD at *t* = 2.21 s and falls back to +0.70 within 0.2 s. The local pairs show nothing of the kind at any point, moving between +0.36 and +0.02 across the same interval. A brief long-range interaction therefore occurs at one moment of a smooth power recovery, with no counterpart in either the power or the local coupling of the sources carrying it.

This is the clearest case in the beta band of long-range coupling without a local counterpart. Inside the window the local pairs stand at +0.20, that is, at their prestimulus level, while the distal estimate is nearly eight times larger. The rank sweep shows the same thing from the other direction: whatever elevation the local estimate has is confined to the lowest ranks, where the projector removes almost nothing, falling from +0.87 prestimulus SD at *R* = 1 to +0.16 by *R* = 200 and fluctuating between +0.01 and +0.17 thereafter, so that it sits on the prestimulus level throughout the regime in which the estimator actually operates. The distal estimate does the opposite, holding between +0.68 and +1.59 across the whole range and peaking at the working rank of 300: it is indifferent to how aggressively the power-dominated subspace is removed. The apparent local coupling of this network is thus almost entirely what the projection takes away, and what it preserves is the long-range interaction alone.

Twenty-nine of the fifty-nine local pairs are nevertheless leakage-resistant, holding between +0.1 and +0.7 prestimulus SD at every rank, while the remaining thirty decay to the prestimulus level, so the cloud of pairs ends slightly above baseline (median +0.16, +0.23 for the resistant subpopulation alone, 49 of 59 pairs above it). The two subpopulations are not distinguishable from one another (median difference +0.041, 95% CI [−0.008, +0.079], *p* = 0.059). A weak residue of local phase alignment therefore accompanies this interaction, an order of magnitude below the long-range effect: IC6 shows long-range coupling that does not depend on local synchrony rather than coupling that excludes it.

IC5 spans 22 sources, linking right inferior parietal and right superior frontal cortex with left superior parietal, precuneus and right paracentral cortex, and peaks at *t* = 1.26 s. Its rise of +6.57 prestimulus SD inside the window is the largest response of the six (Table 2). Unlike IC6 it peaks during the desynchronization itself: the power of its sources reaches a minimum of −3.07 prestimulus SD at *t* = 0.51 s and is still 1.03 SD below baseline when the network peaks.

Its latency places it close to the transient components, and the two must not be confused. IC1 to IC4 peak between 0.81 and 1.01 s, a quarter of a second before IC5, and the raw time course of its pairs shows both: a first rise to +1.35 prestimulus SD near 0.9 s, a return below baseline to −1.25 at 1.06 s, and only then the peak of the network itself. Removing the contribution of the four transients eliminates the earlier rise and leaves the peak standing at +5.66 instead of +6.57, while the local estimate in the window changes from +0.67 to +0.52. All values below refer to the corrected data.

IC5 has the largest local residue of the six, but the comparison that matters is with its own distal estimate. In the window the local pairs stand at +0.52 prestimulus SD against +5.66 for the distal pairs, and at the moment of the peak at +0.49 against +6.65: an order of magnitude apart. The local estimate does not vanish across ranks — 30 of its 34 pairs settle slightly above the prestimulus level, with a median of +0.62, the highest value in either band — but the projection suppresses it far more than it suppresses the distal estimate. The two run close together only at the mildest ranks (local +1.82 against distal +2.05 at *R* = 50); once the projector becomes effective the local estimate falls to between +0.27 and +1.23 while the distal estimate settles between +3.1 and +3.7, a ratio of roughly one to five for the remainder of the range. The split into leakage-resistant and leakage-driven subpopulations adds nothing here (16 against 18 pairs, median difference +0.021, *p* = 0.22), since both lie above baseline rather than on either side of it.

IC5 is therefore not a case of long-range and local coupling rising together in the way alpha IC4 is. It shows a strong long-range interaction, leakage-resistant across the whole range of ranks and occurring while the power of its sources is still more than a standard deviation below baseline, accompanied by a weak but genuine locally coherent residue.

Appendix E reports a deliberately unfavourable variant of the analysis, in which the subspace carrying the movement-related power modulation is projected out before the coupling is estimated and only the four subjects whose desynchronization survives that projection are retained. Distal networks remain recoverable in both bands under those conditions, in windows comparable to those described above.

#### 3.2.4. Phase alignment of the detected interactions

Whether a network is more phase-consistent when it is active is not a question separate from how strongly it is coupled: the coupling factorises exactly into the size of the single-trial cross-terms and the agreement of their phases (Section 2.6). Fig. 12 shows both factors for the six networks, and they behave differently.

**Figure 11:**
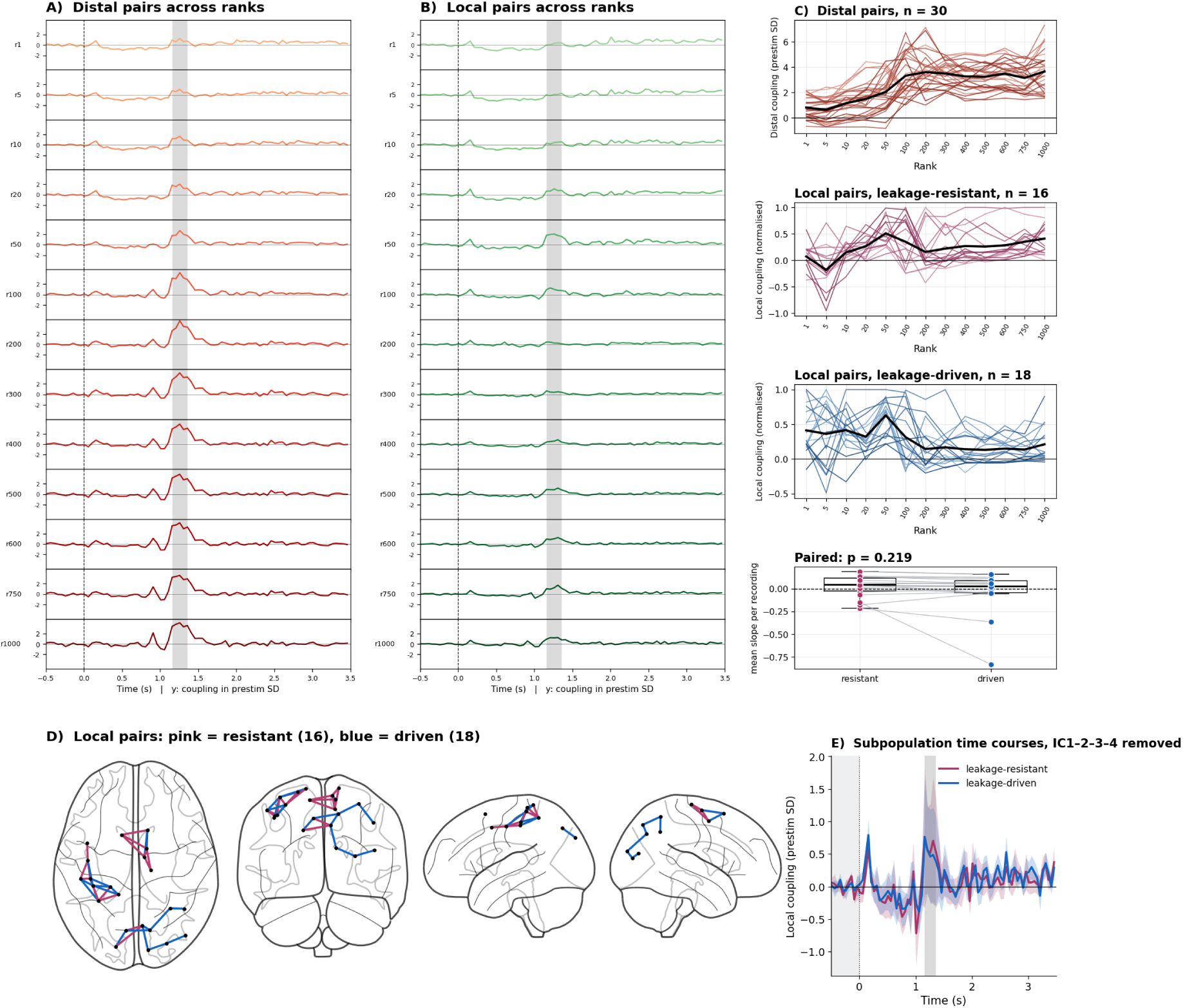
Rank dependence of the local pairs of beta component IC5. As in Fig. 9, without the cortical map of panel D; the raw time courses are in Fig. 10C.

**Figure 12:**
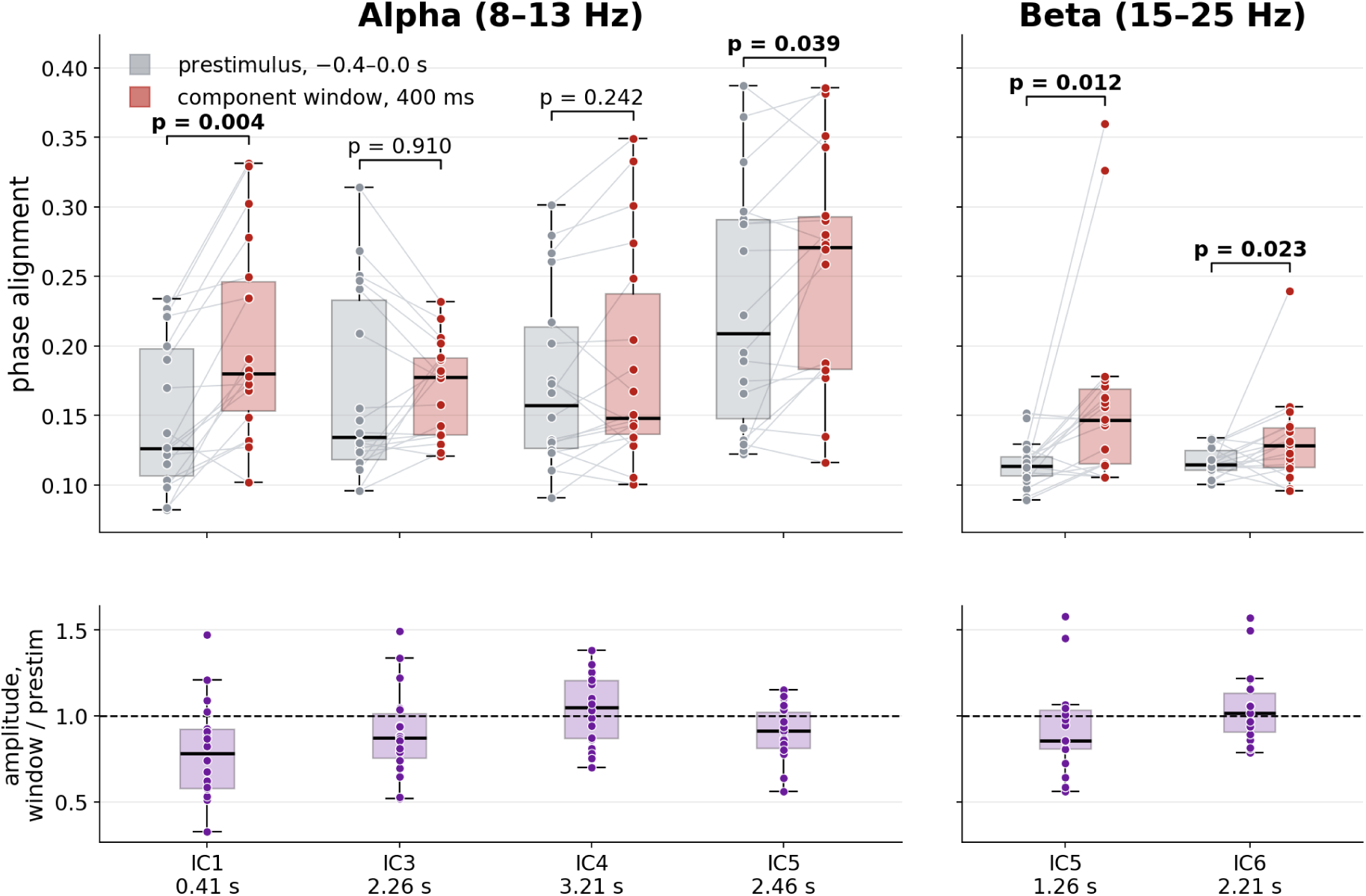
Phase alignment of the single-trial leakage-free cross-terms. Top: the alignment of each component’s 30 pairs in a 400 ms prestimulus interval (−0.4 to 0 s, grey) and in a 400 ms window centred on the component’s own window (red). One point per recording, connected within recordings; *p* is two-sided from exhaustive sign flips at the level of the nine subjects. Bottom: the amplitude factor of the same decomposition, as the ratio of the window to the prestimulus interval, with the dashed line marking equality. A network that lies above its grey box in the top panel and below the dashed line in the bottom panel is one whose coupling is sustained by trials agreeing on a phase relation while the cross-terms themselves shrink.

The early alpha network IC1 raises its phase alignment from 0.149 before the stimulus to 0.204 inside its window (*p* = 0.0039), while the amplitude of its single-trial cross-terms drops to 0.78 of the prestimulus level. The coupling in that window is therefore not held up by larger cross-terms at all — it is held up by the trials converging on a common phase relation while the activity generating them is getting weaker. This is the same window in which the sources of IC1 are most strongly desynchronised, so the two factors move in opposite directions there, and the interaction survives on the one that has nothing to do with power.

The beta network IC5 behaves in a similar way: the prestimulus alignment 0.115 increases up to 0.162 (*p* = 0.012) against an amplitude ratio of 0.86. Alpha IC5 shifts in the same direction, from 0.229 to 0.254 (*p* = 0.039), with its amplitude at 0.91. In both networks the coupling inside the window is supported by the phases of the individual trials agreeing better, and in both the raw coupling stays at roughly its prestimulus level (1.01 and 0.89) even though the cross-terms have shrunk.

Beta IC6 also raises its alignment, from 0.116 to 0.132 (*p* = 0.023), while the amplitude factor is essentially unchanged (1.02) and the raw coupling reaches 1.25 times its prestimulus level. The increase in coupling is thus carried almost entirely by the phase side of the decomposition, without the drop in cross-term amplitude seen in the three networks above.

The two remaining networks make the decomposition worth having, because they show that this is not what always happens. Alpha IC4 does not change its alignment (+0.014, *p* = 0.24), yet it has an amplitude ratio above unity (1.05) and the largest rise in raw coupling of the six (1.27 times the prestimulus level): its window is a genuine growth of the leakage-free cross-terms, not an improvement in their phase agreement. Alpha IC3 moves on neither factor (+0.004, *p* = 0.91).

Four of the six networks show higher phase alignment in their window than in the prestimulus interval, and in three of them — alpha IC1, alpha IC5 and beta IC5 — the amplitude of the single-trial cross-terms is lower at the same time. In these three networks the coupling cannot be attributed to the two regions becoming more active and may indicate a distal phase relation that the individual trials continue to share while the activity carrying it declines.

## 4. Discussion

The present study demonstrates that source-reconstructed MEG, combined with leakage-aware connectivity estimation, can resolve the coexistence of local desynchronization and distal phase coupling within the same cortical nodes. Using controlled simulations and empirical data from a center-out reaching task, we showed that the CD-PSIICOS framework (Kleeva and Ossadtchi, 2025; Altukhov et al., 2023; Ossadtchi et al., 2018) can disentangle these two regimes across a range of projection ranks, frequency bands, and post-movement time windows. The central methodological result is that distal phase coupling remains detectable noninvasively in regions undergoing pronounced event-related desynchronization, including coupling at near-zero phase lag. Conventional estimators cannot resolve this regime: they confuse drops in local power with changes in interareal interaction. This limitation is compounded by the widespread practice of excluding zero-phase-lag interactions to mitigate spatial leakage: Mehra et al. (2025) recently demonstrated that most cortico-cortical functional connectivity occurs at zero or near-zero phase delay, and that discarding these interactions — as done by the imaginary part of coherency, phase-lag index, and related measures — reduces reliability and neurobiological validity of the connectivity estimates. CD-PSIICOS takes advantage of the spatial sampling. It operates in the product space of sensor-space covariance matrices and mitigates spatial leakage through the subspace projection rather than phase-lag exclusion, preserving sensitivity to the full range of physiologically plausible phase relationships within the brain’s functional networks.

Connectivity estimates that stay valid under local power loss matter beyond the specific question addressed here. Recent MEG work shows that distributed connectivity patterns carry cognitively meaningful information that is not reducible to local activation: imagined object categories can be decoded from the spatial covariance between regions, through both short- and long-range connections (Mantegna et al., 2025), and frequency-specific resting-state connectomes of category-selective areas predict individual task performance (Soyuhos et al., 2026). The informational content of such patterns can only be exploited if connectivity estimates remain sensitive when local power decreases, which is the regime the present approach is designed to handle.

The simulation results established detectability: when local phase coherence within two cortical patches was progressively disrupted while distal phase relationships were maintained at the subpopulation level, CD-PSIICOS successfully recovered the distal coupling even as local coupling and spectral power declined. In the empirical MEG recordings, the group decomposition returned six long-range networks, four in the alpha band and two in the beta band, each active in its own time window. They differ in which dynamics survives on the local connections between their own nodes once the power-dominated subspace has been projected out. In one alpha network the leakage-resistant pairs remain below their pre-stimulus level at every rank at which the projector is active, so no stable locally synchronized subpopulation can be identified; in another, both subpopulations settle above that level and rise together with the long-range estimate; the remaining networks lie between these extremes, retaining a residue that is real but an order of magnitude weaker than the long-range coupling of the same sources. The two extremes are also the extremes of concurrent power suppression: the network with no local counterpart peaks while the power of its sources is below baseline, whereas the one in which local and long-range coupling rise together peaks after power has returned to its pre-stimulus level. The simulations do not establish by which mechanism this gradient of regimes arises in the brain; the paired-subpopulation model that generated them is one candidate, and we consider it alongside alternative accounts below.

The earliest alpha network, in which distal coupling rises with no strong local oscillatory support, is most naturally accommodated by frameworks that do not require local synchrony as a prerequisite for interareal coordination. The Synaptic Source Mixing model (Schneider et al., 2021) proposes that field-field coherence arises as a byproduct of spiking transmission through anatomical connections: postsynaptic potentials generated in a target region by afferent spikes produce interareal coherence without requiring the target to be locally oscillating. Under this interpretation, the alpha-band distal coupling detected by CD-PSIICOS may index the activation state of cortico-cortical pathways rather than the local oscillatory state of either endpoint. Related frameworks — Communication Through Resonance (Hahn et al., 2014), transient synchrony routing (Palmigiano et al., 2017), and coherence-through-communication (Vinck et al., 2023) — similarly predict that interareal phase coordination can arise without a sustained local oscillatory synchrony.

An alternative, though not mutually exclusive, interpretation draws on the subpopulation-level account. Theoretical work on chimera states (Kuramoto and Battogtokh, 2002; Abrams and Strogatz, 2004; Wang and Liu, 2020; Dogonasheva et al., 2025) and heterogeneous neural networks (Dahmen et al., 2022, 2019) demonstrates that macroscopic desynchronization is compatible with coherent phase relationships maintained by subsets of neurons. The MEG signal, reflecting the aggregate activity of large neural populations, may average over these subpopulation dynamics, registering ERD while coherent subgroups sustain interareal phase alignment. In the balanced excitation–inhibition regime (Brunel, 2000; Renart et al., 2010), near-zero pairwise spiking correlations coexist with population-level oscillatory structure — a direct dissociation between cellular synchrony and macroscopic phase organization that could support the patterns we observe.

Consistent with this account, recent OPM-MEG recordings during motor imagery have shown that the beta-band spectral peak (12–25 Hz) persists throughout desynchronization, with movement-related modulations manifesting primarily as reductions in peak amplitude rather than abolition of the oscillation itself (Fedosov et al., 2025). This suggests that macroscopic ERD is compatible with the continued operation of more spatially restricted oscillating clusters whose aggregate contribution survives at the sensor level and could plausibly scaffold the distal coupling detected here.

An independent line of evidence comes from the decomposition of the coupling itself. Because the projector is linear and commutes with averaging over trials, the reported coupling factorises exactly into the amplitude of the single-trial cross-terms and the consistency of their phases, and the second factor is normalised by the first: it cannot increase merely because the two regions have become more active. Four of the six networks are more phase-consistent inside their window than before the stimulus, and in three of these the amplitude of the cross-terms is lower at the same time. The coupling in those windows is therefore sustained by individual trials converging on a common phase relation while the activity carrying it declines, which is what the subpopulation-level account predicts and what a residual power effect cannot produce. The exception is instructive. The one network whose local pairs rise above baseline together with its long-range estimate is also the only one whose phase consistency does not change; there the increase in coupling is a growth of the leakage-free cross-terms themselves. The two analyses thus separate the same cases from opposite directions, one through the rank dependence of the short connections and the other through the trial-to-trial phase structure of the long ones.

A further alternative is that local and distal coupling reflect two physically distinct oscillatory channels coexisting at the same cortical site – an intrinsic microcircuit oscillation that desynchronises with movement, and a region-wide carrier whose phase is paced by a separate generator and remains locked to its counterpart in the partner region. A plausible substrate would assign the local channel to supragranular pyramidal– interneuron loops and the distal channel to long-range projection neurons gated by a higher-order thalamic relay such as the pulvinar (Saalmann et al., 2012; Halgren et al., 2019). Under this account, distal coupling can remain stable even when both endpoints are locally desynchronised because the carrier of the distal channel has its own oscillatory source, anatomically distinct from the microcircuit that produces the desynchronization.

The two beta networks illustrate the two interpretations side by side. The earlier one peaks while the power of its sources is still more than a standard deviation below baseline, and its long-range coupling is accompanied by a genuine local residue: most of its short connections settle above their pre-stimulus level once the power subspace has been removed, although the coupling they carry is several times smaller than the distal estimate. A subpopulation-mediated mechanism would produce exactly this: a spatially restricted subset retaining local phase coherence sufficient to support the distal interaction while the bulk population is desynchronized. The later network behaves differently. Its short connections sit at their pre-stimulus level throughout the range of ranks in which the estimator operates, and its coupling event has no counterpart either in local coupling or in the power of its sources, which recovers smoothly through the same interval without an excursion of its own. A brief long-range interaction occurring against an unmodulated local background is more readily accommodated by an account in which the distal channel has its own generator, anatomically distinct from the microcircuit that produces the desynchronization.

### 4.1. Limitations

The generative model of the simulations does not include burst-like or non-stationary dynamics, which contribute to the dissociation between phase- and power-based networks (Hindriks and Tewarie, 2023); the empirical coupling estimates are averaged over trials and over 200 ms windows and reflect the mean phase relation within a window rather than coupling resolved at the level of individual bursts.

### 4.2. Perspectives

The methodological framework demonstrated in this paper provides a general tool for investigating the local–distal coupling relationship from noninvasive recordings. Several extensions are warranted. Simultaneous intracranial–MEG recordings would allow direct validation of the subpopulation-level interpretation by comparing MEG-derived coupling estimates with local field potential and single-unit measures of phase organization. Application to clinical populations with altered oscillatory dynamics — such as Parkinson’s disease, where beta synchronization is pathologically enhanced — could test whether the dissociation between local and distal coupling is disrupted under conditions of excessive local synchrony. Finally, resting-state analyses could establish whether the observed regimes reflect task-specific network reconfiguration or a more general property of cortical dynamics.

## 5. Conclusion

This study demonstrates that MEG, equipped with leakage-aware connectivity estimation, can detect and disentangle two distinct coupling regimes: one in which distal phase coordination coexists with macroscopic local desynchronization, and one in which both operate together. These results establish that local synchrony is not a necessary prerequisite for long-range phase coupling at the macroscopic level and that this dissociation is accessible to noninvasive neuroimaging when appropriate methodological tools are employed.

## 7. Data and Code Availability

The empirical MEG data analyzed in this study are openly available from (Yeom et al., 2023). The CD-PSIICOS framework is implemented in an open-source Python package available at https://github.com/dkleeva/CD-PSIICOS.

## 9. Ethics approval

The original data collection (Yeom et al., 2023) was approved by the Institutional Review Board of Seoul National University Hospital (approval number 1105-095-363), with written informed consent from all participants. The present study is a secondary analysis of openly available de-identified data and did not require additional ethical approval. All procedures complied with the Declaration of Helsinki and with relevant laws and institutional guidelines.

## 10. Appendix A. The simulation benchmarks for projection rank and the spatial profile of the coupling estimate

Figure 13 illustrates a single Monte-Carlo realisation of the generative model. The figure summarises three complementary sensor-level observables across the full within-region jitter sweep *σ*_local_*/π* ∈ [0, 2]. At *σ*_local_ = 0 the two ROIs project a clear focal power pattern onto the planar gradiometers (Fig. 13B, leftmost panel). As within-region jitter grows, the pattern attenuates rapidly: by *σ*_local_ = 0.3*π* the focal blob has collapsed by roughly half, and by *σ*_local_ ≥ 0.5*π* the topography becomes essentially indistinguishable from background. Critically, no parameter of the source signal has changed: each cortical vertex continues to emit a sinusoid at the same fixed amplitude *A* across the entire sweep. The collapse arises purely from destructive interference at the macroscopic ROI level once subpopulation phases desynchronise.

**Figure 13:**
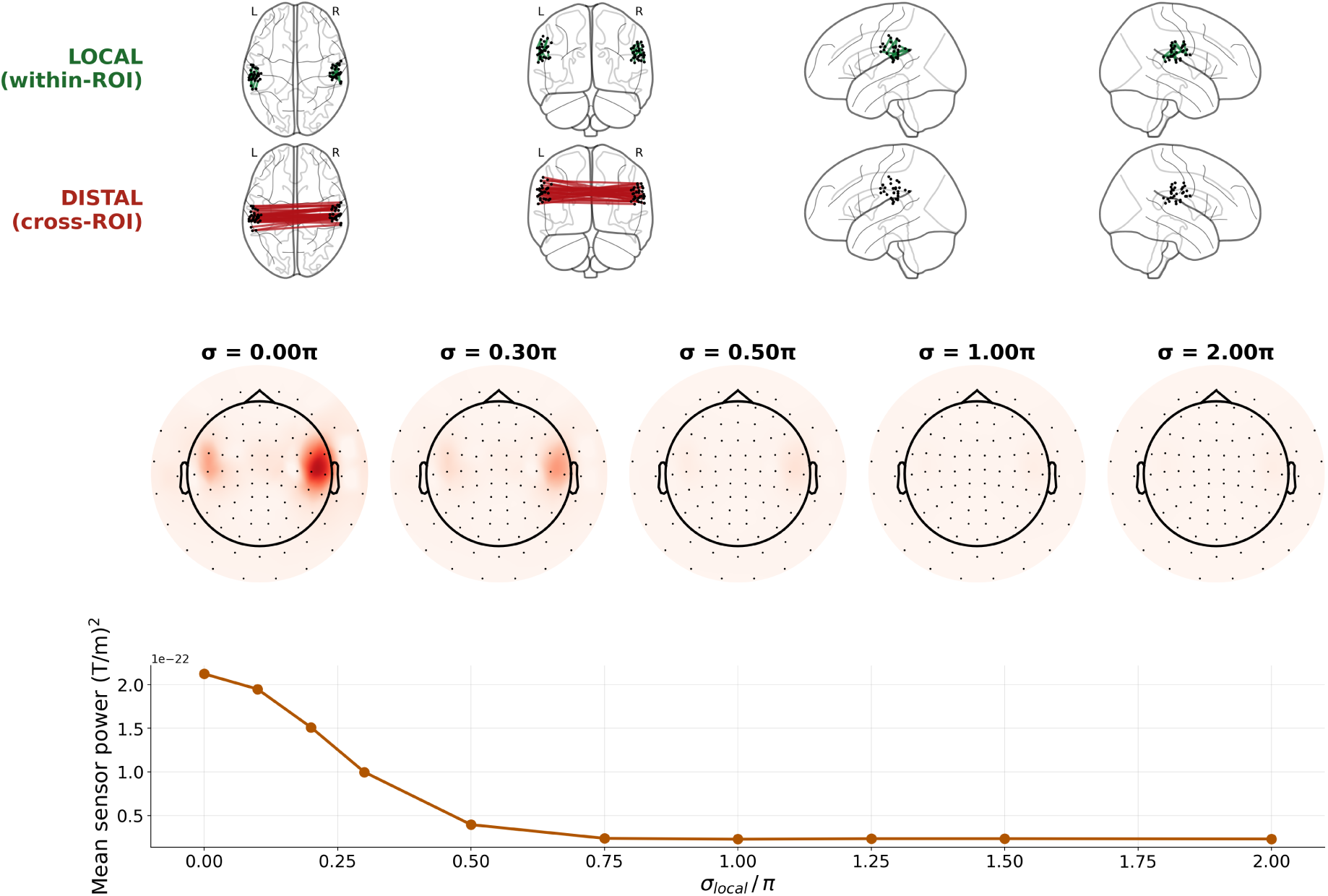
Representative simulation example (*K* = 30, Δ = *π/*20, *f*_0_ = 10 Hz, single MC).(A) Ground-truth networks on an MNI glass brain: local within-ROI edges (green, top), distal paired cross-ROI edges *L_k_* ↔ *R_k_*(red, bottom). (B) Sensor-power topographies at *σ*_local_*/π* ∈ {0, 0.3, 0.5, 1, 2}, shared colour scale. (C) Mean sensor power vs *σ*_local_*/π*.

The mean sensor power (Fig. 13C) drops by roughly an order of magnitude over the range *σ*_local_*/π* ∈ [0, 0.75], then saturates at a noise floor for the remainder of the sweep (*σ*_local_ ≥ *π*). The location of the saturation knee at *σ*_local_ ≈ 0.5*π* is diagnostic: it corresponds to within-ROI phase dispersion comparable to the radian unit, beyond which the coherent macroscopic sum is already swamped by destructive cancellation and the visible MEG signature reduces to the residual contribution of the distal coupling channel plus instrumental noise.

The projection rank *R* controls how much of the power-dominated subspace is removed, and therefore how far the estimate is pulled away from the local connections on which spatial leakage persists.

To establish the operating regime of the projection rank *R*, we applied the full pipeline at *σ*_local_ = 0 (i.e. all subpopulations in phase, the condition under which both local leakage and genuine distal coupling are simultaneously maximal) and inspected how pair-wise coupling varies with inter-source distance for each *R* ∈ {0, 1, 5, 10, 20, 50, 100, 300, 500, 700} (Fig. 14). Per-rank curves were normalized to the peak distance bin to make their shape directly comparable.

**Figure 14:**
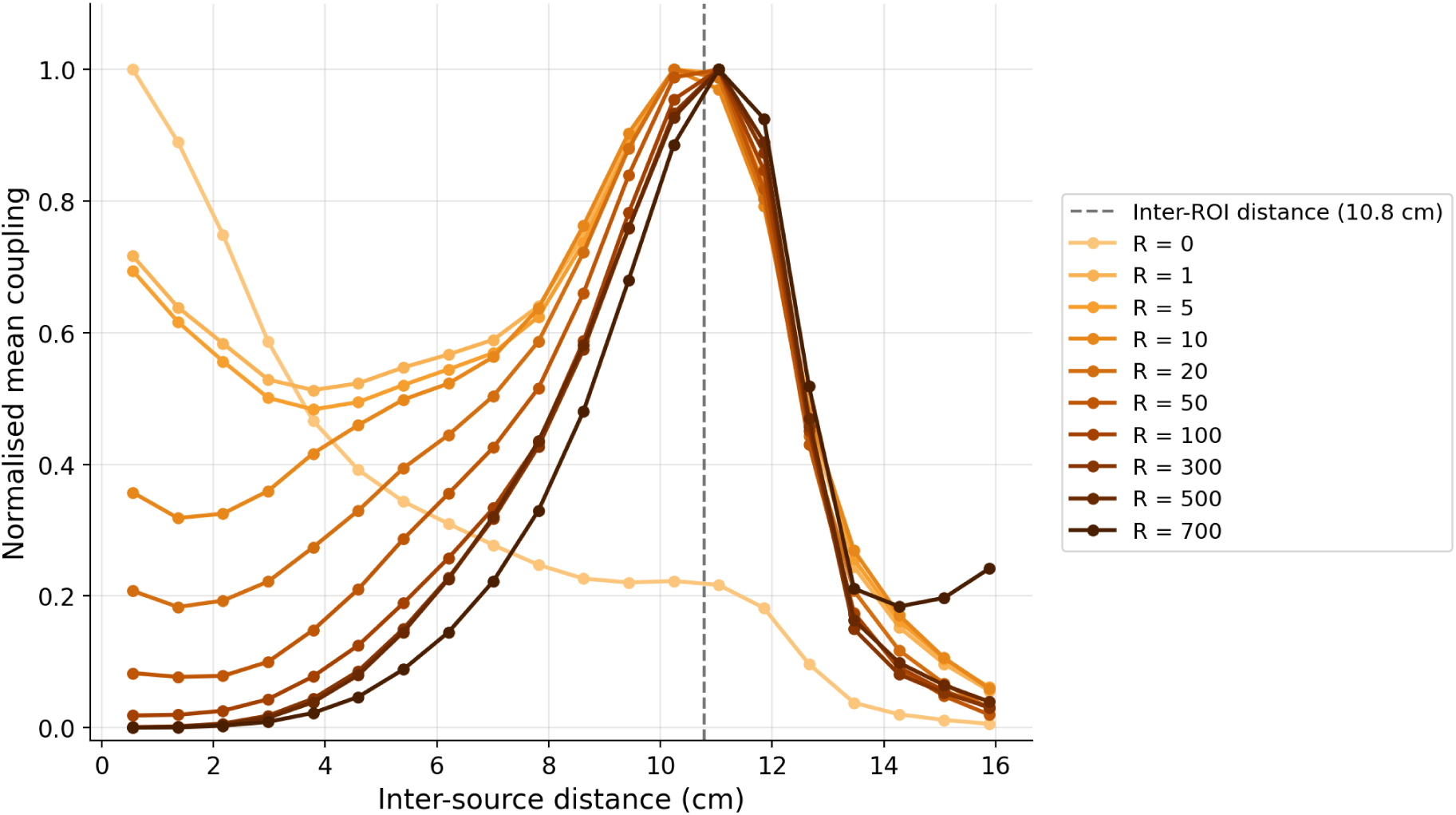
Distance dependence of pair-wise CD-PSIICOS coupling computed from the simulated data at *σ*_local_ = 0 across projection ranks *R* observed. For each rank the coupling magnitude was computed for all source pairs, binned by inter-source Euclidean distance, and normalized to the peak bin to compare curve shapes across ranks. The dashed grey line marks the geometric inter-ROI centroid separation (10.8 cm)

**Figure 15:**
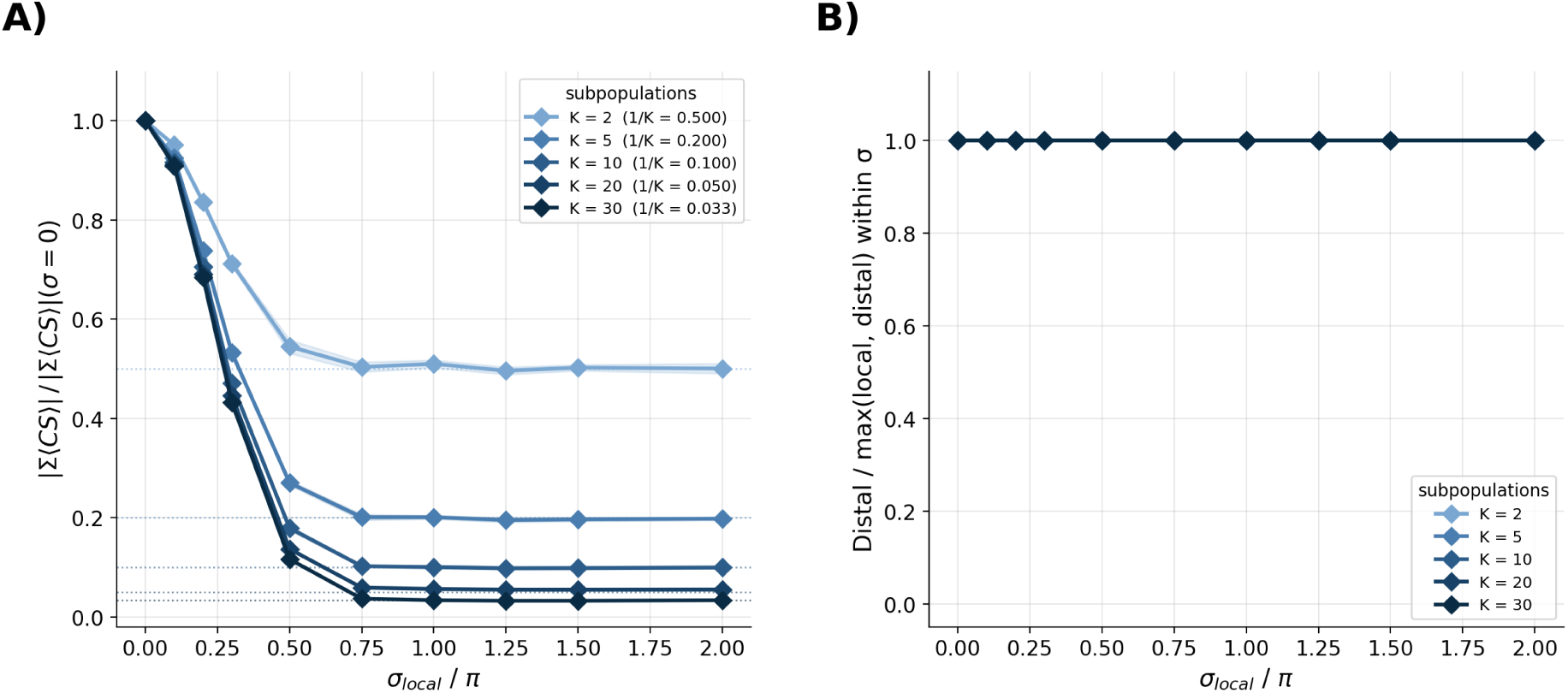
Distal coupling across subpopulation counts *K* ∈ {2, 5, 10, 20, 30}. (A) Source-level distal coherent sum *S*^dist^ normalized to its *σ* = 0 value. Each curve saturates at the theoretical 1*/K* plateau (dotted reference lines): 0.50 for *K*=2, 0.20 for *K*=5, 0.10 for *K*=10, 0.05 for *K*=20, 0.033 for *K*=30. (B) The same statistic normalized within each *σ* by the maximum of the local and distal coherent sums

At rank *R* = 0 (no projection, lightest curve) the global maximum of the distance profile sits at the nearest-neighbour bin — the canonical signature of spatial leakage between sources with overlapping leadfields — but a clear secondary peak is already present at the 10.8 cm inter-ROI distance, reaching ≈ 0.45 of the leakage maximum. At *R* = 1 the situation reverses: a single-rank projection is sufficient to suppress the short-range leakage component to roughly half its previous level, and the distal peak at 10.8 cm becomes the new global maximum. Increasing *R* further continues to push down the residual short-range contribution while the distal peak sharpens, so that by *R* ∈ {100, 300, 500, 700} (dark orange) the curves are essentially zero at small distances and exhibit a sharp, well-localised peak at the ground-truth inter-ROI centroid distance (dashed grey line), recovered to within one distance bin.

This analysis confirms that CD-PSIICOS faithfully recovers the spatial scale of long-range interactions: the peak landing exactly at the geometric inter-ROI distance is direct evidence that the surviving cross-spectrum is interaction-specific rather than numerical noise. Second, the broad rank plateau over which the peak location is stable (*R* ∈ [1, 700]) shows that the choice of projection rank is forgiving with respect to spatial fidelity. Combined, these observations license the use of *R* = 100 in the analyses for simulations described in the main text.

Two theoretical properties follow from the paired-jitter construction, derived from the generative model and confirmed at the source level.

- *Within-ROI coupling collapses.* The cross-spectrum between two distinct vertices(*i, j*) of the same ROI averages to ⟨*C_ij_*⟩ = *A*^2^ exp(−σ^2^_local_) when *s*(*i*) ≠ *s*(*j*), and to *A*^2^ only when *s*(*i*) = *s*(*j*). With *K* = *N_v_* (the *K* = 30 regime used throughout) every within-ROI pair has distinct subpopulation labels, so mean within-ROI coupling decays to zero as *σ*_local_ → ∞.
- Distal coupling reaches a 1/K plateau. The cross-ROI coherent sum equals

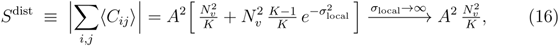

so between-ROI pair coupling saturates at a non-zero plateau determined by *K*, the number of paired subpopulations.

Note that only cross-region pairs sharing a subpopulation index are paired; the remaining fraction 1 - 1/K of cross-region pairs receive independent jitter and decay as exp(−σ^2^_local_), exactly as within-region pairs. The limit *K* → ∞ therefore corresponds to a fully unpaired model in which no distal coupling survives within-region desynchronization. Since both regions share a common per-epoch base phase, the simulated distal coupling is equivalent to a common drive with a fixed lag; the estimator quantifies the consistency of the inter-regional phase relation and does not distinguish direct interaction from shared input.

To test how the assumed microarchitecture of populations influences the recovery of long-range coupling under within-region jitter, we swept the subpopulation count over *K* ∈ {2, 5, 10, 20, 30} at fixed inter-region phase lag Δ = *π/*20, and quantified the source-level distal coherent sum *S*^dist^ ≡ |∑*_ij_*_∈_*_L_*_×_*_R_*⟨*C_ij_*⟩| for every value of *σ*_local_ ∈ [0, 2*π*]. Two complementary normalizations of *S*^dist^ are shown in Figure 15.

When each curve is normalized to its own value at *σ*_local_ = 0, every *K* reproduces a smooth decay that saturates by *σ*_local_ ≈ *π* at a plateau that lies almost exactly on the theoretical 1*/K* horizontal reference line (dotted): 0.50 for *K*=2, 0.20 for *K*=5, 0.10 for *K*=10, 0.05 for *K*=20 and 0.033 for *K*=30. This confirms that the model behaves as predicted analytically — the saturation level of *S*^dist^ is set by the fraction of cross-region pairs that share their per-epoch phase jitter ((*N_v_/K*)^2^ · *K = N^2^_v_/K* paired pairs out of *N^2^_v_*). The fraction of distal coupling that survives within-region jitter is therefore an inverse function of how finely the cortex is subdivided into independent oscillatory units, ranging from one half in the most coarsely clustered regime to a few percent in the more vertex-resolved regime.

The decay shown in the left panel might at first suggest that distal coupling becomes *undetectable* relative to local coupling at large *σ*_local_. The right panel, where each distal trace is divided by the larger of the local and distal coherent sums at the same *σ*_local_, shows that this is not the case. Across all *K* the within-*σ*-normalized distal trace stays at unity — the distal coherent sum is the maximum of the two pair classes at every jitter level. The reason is straightforward: while the absolute distal sum decays from ∝ *N* ^2^ to ∝ *N_v_/K*, the local coherent sum collapses entirely toward zero as soon as within-region phase coherence is destroyed, so the distal-to-local ratio actually grows with *σ*_local_, not shrinks. The result holds independently of *K*: even at *K* = 30, where only ≈ 3.3% of the original distal magnitude remains at *σ*_local_ ≥ *π*, that 3.3% is still substantially larger than the within-ROI residual.

## 11. Appendix B. Limitations of imaginary coherence

The simulations reported in Section 3.1 were performed at a single inter-regional phase lag Δ = *π/*20, slightly offset from zero. This choice is not innocent: imaginary part of cross-spectrum (denoted as ImCS later) is, by construction, blind to zero-lag interactions. Although this measure is therefore commonly framed as a leakage-robust estimator, the same property that protects it against zero-lag spatial leakage also forces it to mis-rank *any* coupling that happens to be near-instantaneous,including, in our generative model, every within-ROI coupling. Local pairs, whose subpopulation phases differ only by the shared jitter *ε_k_* and whose mean phase difference is therefore zero, are systematically suppressed by ImCS. Cross-ROI pairs, whose phases differ by Δ = *π/*20 ≠ 0, retain a non-trivial imaginary component and survive. This effect is that ImCS is structurally biased *toward* the distal class in our benchmark. This bias makes ImCS appear competitive with CD-PSIICOS; it is also exactly what fails when the distal lag itself collapses toward zero, as the rest of this subsection makes clear.

We expose this dependency with two complementary analyses run on identical simulated data: we compared the performance with regard to network reconstruction at two contrasting distal phase lags (Fig. 16), and a continuous phase- and jitter-lag sweep (Fig. 17). Per-pair detection performance is summarised by two complementary curves. The receiver operating characteristic (ROC) plots the true-positive rate (TPR) against the false-positive rate (FPR) as the threshold is swept across the range of cross-spectral coefficients over all pairs. The precision-recall (PR) curve plots precision (positive predictive value) against recall (TPR) under the same sweep, and its area (PR-AUC, equivalent here to the average precision) is the classifier’s mean precision across all recall levels. ROC-AUC is the natural global summary, but on this benchmark it is dominated by the high-FPR regime; because the ground-truth set of distal pairs is small relative to the full pair list, PR-AUC is the more sensitive metric in the high-specificity corner where method differences are observed, and we therefore report both throughout.

**Figure 16:**
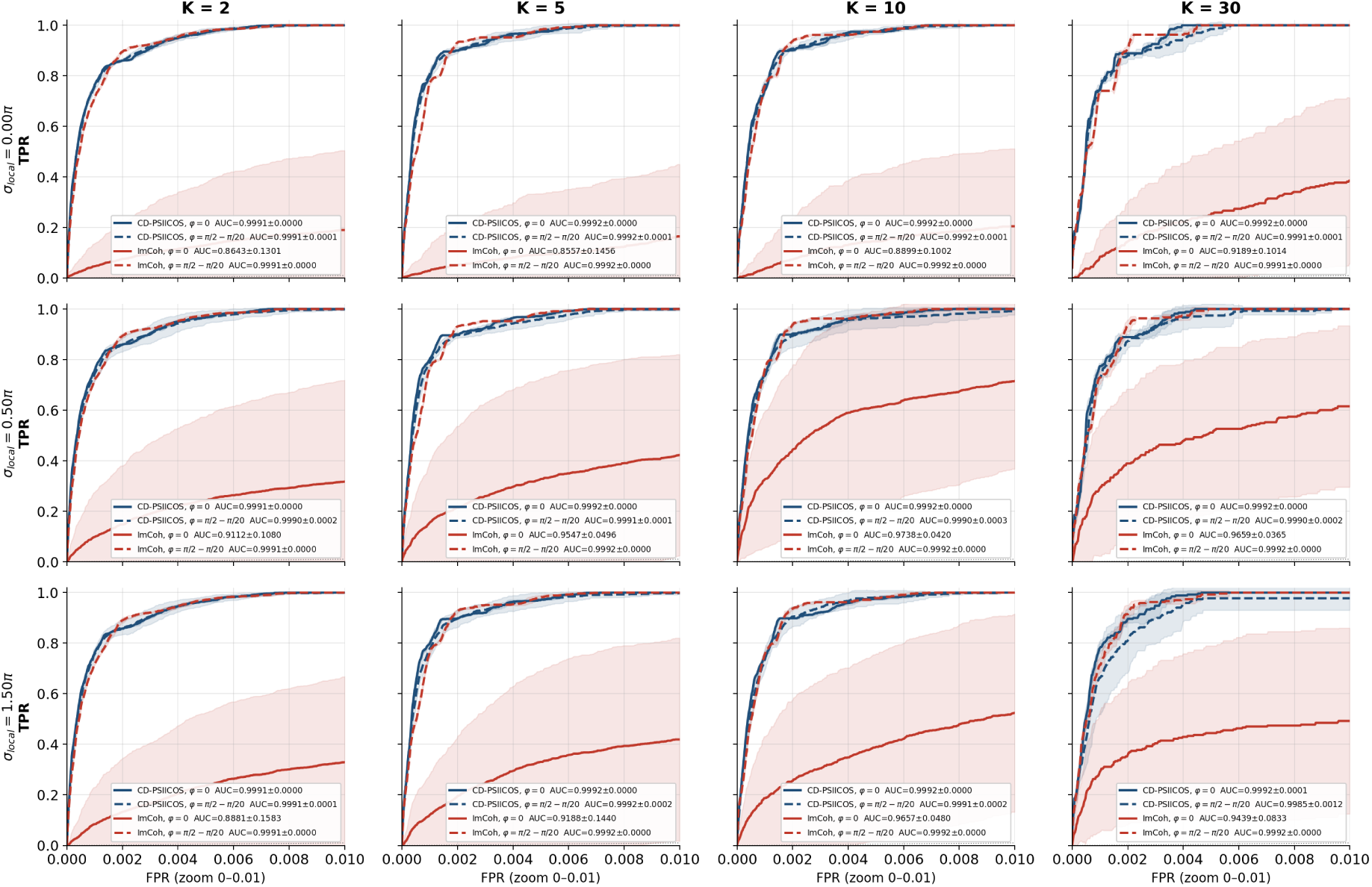
Per-pair detection ROC for CD-PSIICOS (blue) and ImCS (red) on identical simulated MEG data, evaluated at two phase-lag scenarios per cell: *δ* = 0 (solid) and *δ* = *π/*2 − *π/*20 (dashed). Rows: within-region jitter *σ*_local_*/π* ∈ {0, 0.5, 1.5}; columns: subpopulation count *K* ∈ {2, 5, 10, 30}

**Figure 17:**
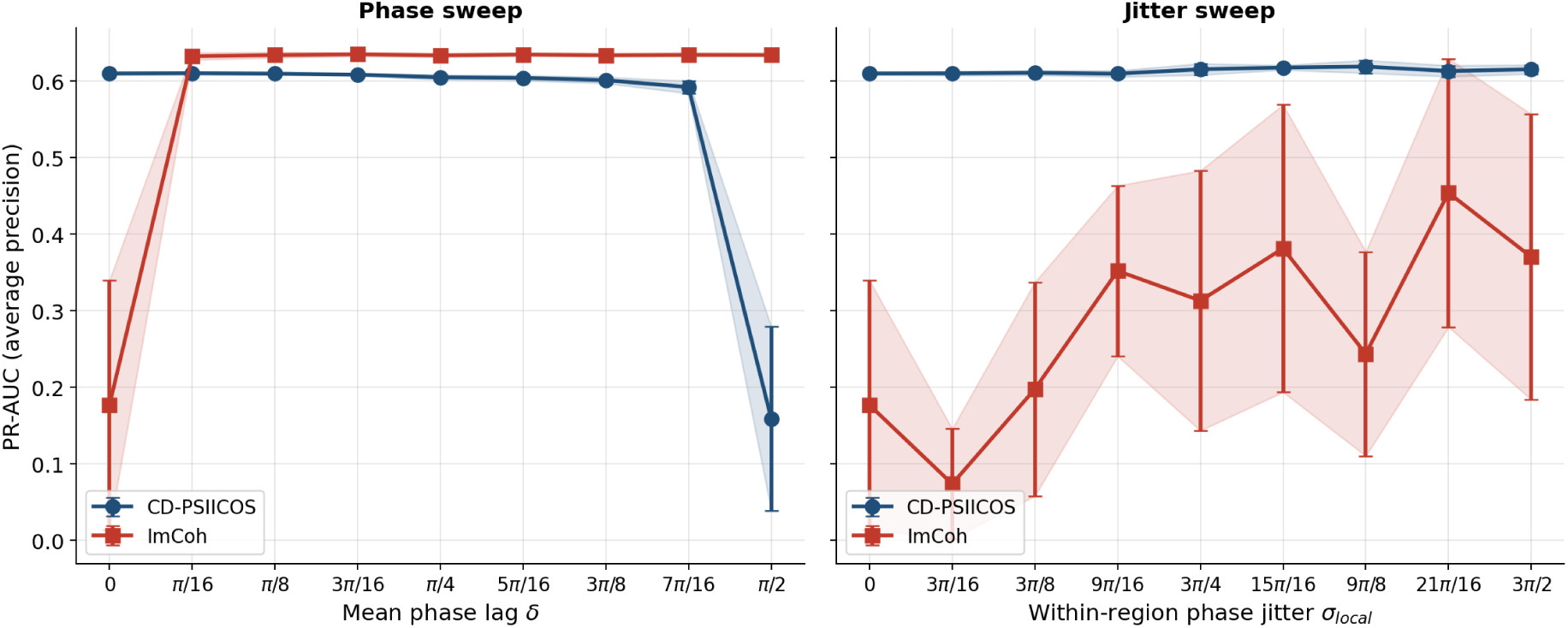
PR-AUC sweeps for CD-PSIICOS (blue) and ImCS (red) at *K* = 30. Left: phase-lag sweep at fixed within-region jitter *σ*_local_ = 0. Right: within-region jitter sweep at fixed inter-regional lag *δ* = 0 (ImCS blind spot). Markers are mean AUC across *N*_MC_ Monte-Carlo trials, shaded bands are ±1*σ* across MC.

Figure 16 reports per-pair detection ROCs. Each cell is computed at two contrasting phase scenarios on the same simulated MEG data: the ImCS-blind condition *δ* = 0, and the CD-PSIICOS adversarial condition *δ* = *π/*2 − *π/*20. Three of the curves (both CD-PSIICOS curves for both lags and the ‘lagged’ ImCS curve ImCS*_δ_*_=*π/*2−*π/*20_) are essentially indistinguishable across the entire set of scenarios, with mean AUCs locked at ≈ 0.9991–0.9992. The only curve that departs from this observation is the zero-lag ImCS curve ImCS*_δ_*_=0_ (red solid), and it does so in every scenario. The mean ROC sags below the others in the high-specificity corner, with mean AUC ranging from 0.86 at (*σ*_local_ = 0*, K* = 2) to 0.97 at(*σ*_local_ = 0.5*π, K* = 10).

The comparison yields two diagnostics. First, the contrast between ImCS*_δ_*_=*π/*2−*π/*20_ (mean AUC 0.9992, std *<* 10^−3^) and ImCS*_δ_*_=0_(mean AUC 0.86–0.97, std up to 0.16) on the same data is a direct diagnostic of the distal-class bias discussed above: ImCS inherits the apparent ceiling performance of the asymmetric scenario from the structural suppression of local zero-lag pairs, and loses it the instant the distal lag also collapses to zero. Second, the dependence of the ImCS*_δ_*_=0_ failure on *K* and *σ*_local_ is non-monotonic: the lowest mean AUCs sit at *σ*_local_ = 0 (where every local pair is fully coherent and therefore maximally suppressed by the imaginary projection, leaving the smallest residual signal-to-noise margin for the surviving distal pairs), and the failure relaxes somewhat as *σ*_local_ grows.

To map the dependence continuously and to expose the precision of each estimator in the high-specificity regime, we evaluated PR-AUC at fixed subpopulations number *K* = 30 along two parameters (Fig. 17). The first axis varies the inter-regional phase lag *δ* ∈ [0*, π/*2] at *σ*_local_ = 0; the second varies the within-region jitter *σ*_local_ ∈ [0, 3*π/*2] at *δ* = 0. The two configurations share the operating point (*δ, σ*_local_) = (0, 0), which by construction returns the same value in both panels. The dependence on *δ* at *σ*_local_ = 0 (Fig. 17, left) reveals a structurally symmetric pair of failure points, one for each estimator. ImCS delivers PR AUC ≈ 0.2 at *δ* = 0, then rises sharply to ≈ 0.63 by *δ* = *π/*16 and holds that plateau across the rest of the range. CD-PSIICOS exhibits the complementary profile: it maintains PR AUC ≈ 0.61 from *δ* = 0 through *δ* = 7*π/*16, then collapses to ≈ 0.2 at *δ* = *π/*2.

**Figure 18:**
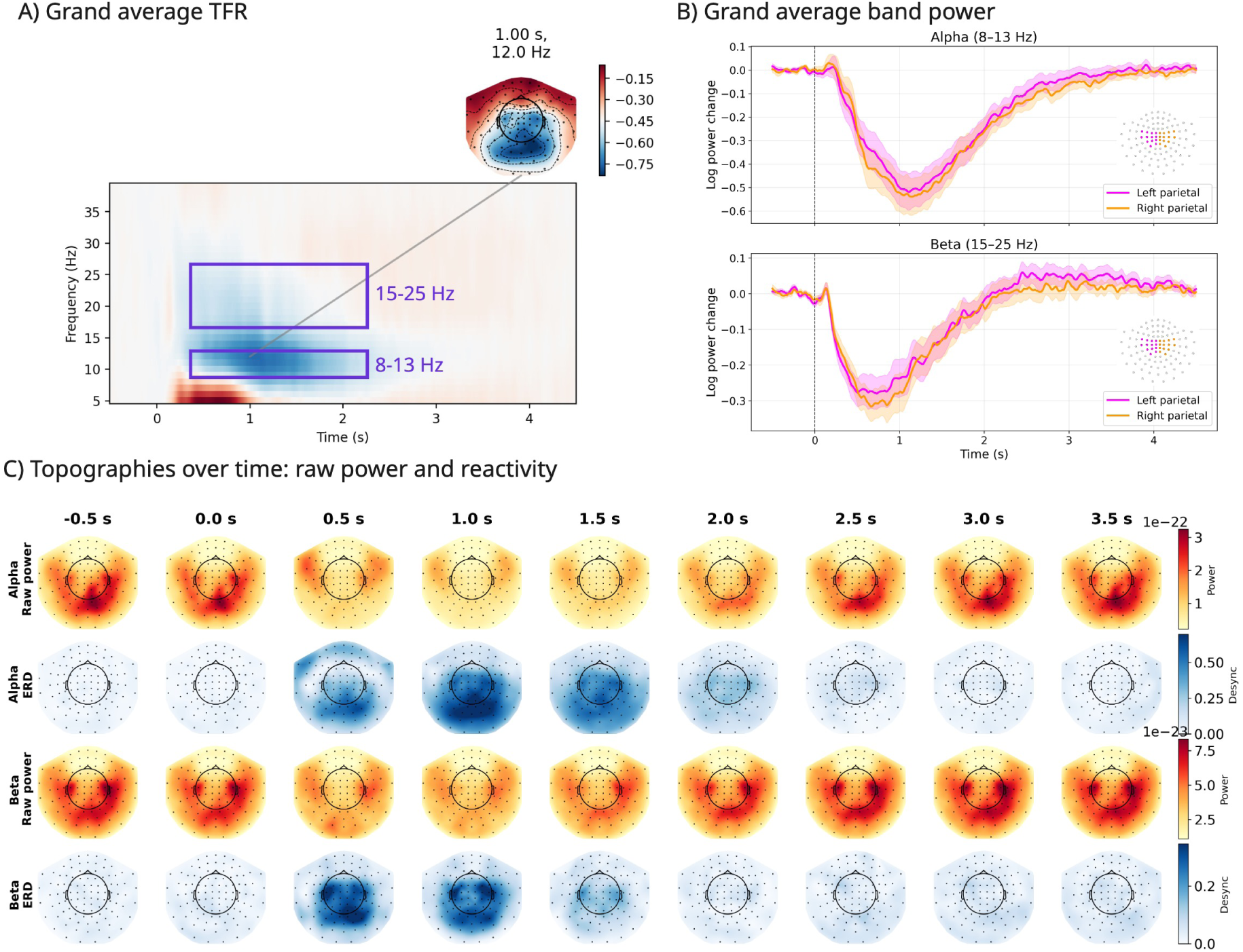
Movement-related time–frequency power modulations in planar gradiometer signals. (A) Grand-average time–frequency representation showing log-ratio baseline-corrected power changes relative to the pre-stimulus interval. Purple rectangles indicate the alpha(8–13 Hz) and beta (15–25 Hz) frequency bands selected for further analysis. (B) Temporal evolution of log-ratio baseline-corrected band power in the alpha (top) and beta (bottom) bands, averaged over left (magenta) and right (orange) parietal sensor groups. Shaded areas indicate variability across subjects and sessions. Vertical dashed lines denote movement onset. (C) Topographic time series showing raw power (rows 1, 3) and baseline-normalized desynchronization index (rows 2, 4) for alpha and beta bands at successive time points. Desynchronization is most pronounced over occipital and sensorimotor regions, particularly in the beta band.

**Figure 19:**
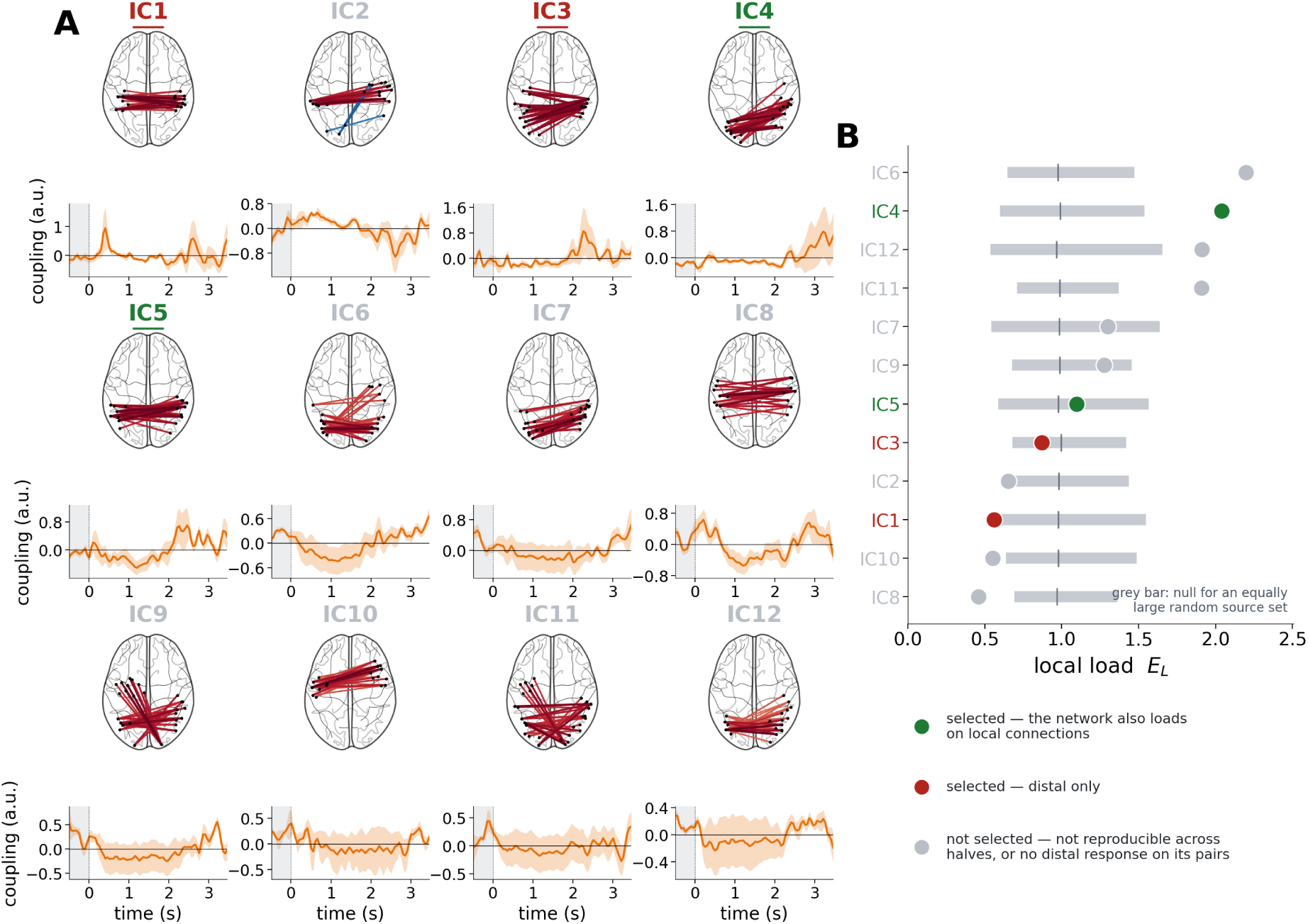
Group decomposition of alpha-band coupling. (A) The twelve temporal-ICA components obtained from the concatenated coupling time courses of all 18 recordings. For each component, the glass brain shows the 30 source pairs carrying the largest absolute mixing weight, with edge colour indicating the sign and magnitude of the weight; the panel below shows the component’s time course as the mean over the nine subjects ± SEM, in arbitrary units. The grey band marks the prestimulus interval and the dotted line the stimulus onset. Component labels are underlined when the component was retained and are coloured as in panel B; components that were not retained are shown in grey. (B) Local load *E_L_* of each component: the mean absolute weight placed on the short connections of the component’s own sources, divided by the mean absolute weight placed on all other short connections. Grey bars give the 5–95% range of the corresponding null, obtained from the random source sets of the same size as the component’s node set, with the tick marking its median. Retained components are coloured green when their local load lies above the null median and red when it lies below; components that were not retained are grey.

**Figure 20:**
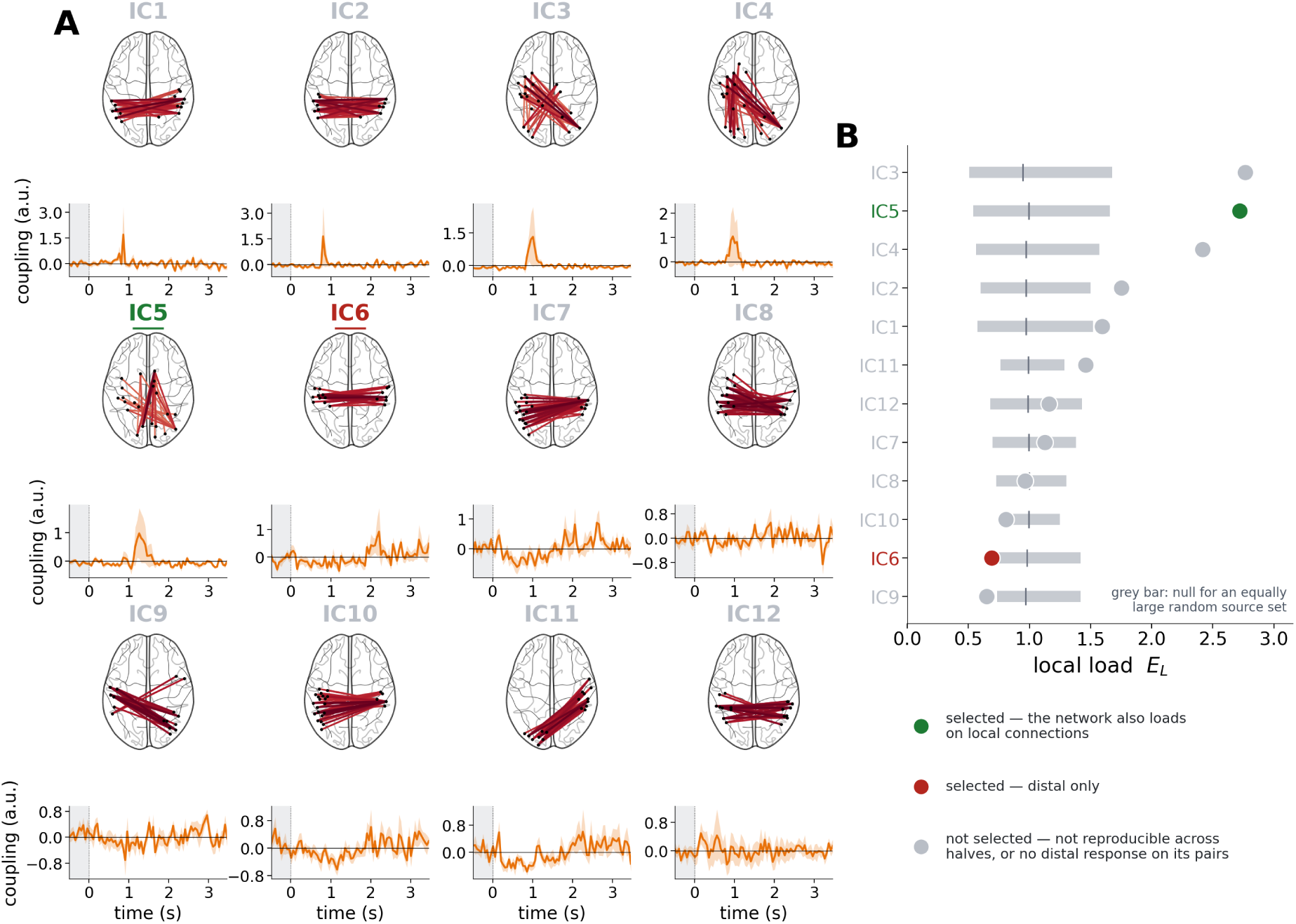
Group decomposition of beta-band coupling. (A) The twelve temporal-ICA components obtained from the concatenated coupling time courses of all 18 recordings. For each component, the glass brain shows the 30 source pairs carrying the largest absolute mixing weight, with edge colour indicating the sign and magnitude of the weight; the panel below shows the component’s time course as the mean over the nine subjects ± SEM, in arbitrary units. The grey band marks the prestimulus interval and the dotted line the stimulus onset. Component labels are underlined when the component was retained and are coloured as in panel B; components that were not retained are shown in grey. (B) Local load *E_L_* of each component: the mean absolute weight placed on the short connections of the component’s own sources, divided by the mean absolute weight placed on all other short connections. Grey bars give the 5–95% range of the corresponding null, obtained from the random source sets of the same size as the component’s node set, with the tick marking its median. Retained components are coloured green when their local load lies above the null median and red when it lies below; components that were not retained are grey

Along the within-region jitter axis at the ImCS-blind operating point *δ* = 0, CD-PSIICOS holds a flat plateau at PR AUC ≈ 0.61, while ImCS starts at PR AUC ≈ 0.2 at *σ*_local_ = 0, dips to a minimum PR AUC ≈ 0.08 at *σ*_local_ = 3*π/*16, and climbs erratically toward ≈ 0.37–0.45 at large *σ*_local_. The climb is not a recovery of sensitivity, as the underlying distal lag is unchanged, but a consequence of within-region jitter producing a small sampling-driven imaginary residual that ImCS partially exploits. ImCS never reaches the CD-PSIICOS plateau.

## 12. Appendix C. Sensor-level characterisation of the movement-related power modulations

Figure 18A shows the grand-average time–frequency representation of the planar gradiometer signals, averaged across subjects, with a pronounced reduction of oscillatory power beginning shortly after movement onset and persisting throughout the movement execution period until the typical rebound onset. The desynchronization was most prominent in the alpha/mu (8–13 Hz) and beta (15–25 Hz) bands, which were selected for all analyses reported in the main text.

Panel B summarizes the temporal evolution of band-limited power averaged over left and right parietal sensor groups. In both frequency bands a clear decrease followed movement onset, reaching a minimum approximately 0.8–1.2 s after onset, followed by a gradual rebound.

Topographic maps computed at representative time–frequency points (panel C) show that movement-related power suppression was not restricted to sensorimotor sensors. In the alpha/mu band desynchronization extended toward posterior sensors, indicating additional involvement of occipital regions; a similar but weaker posterior contribution was also observable in the beta band. This distribution is consistent with the source-space power maps reported in Section 3.2.1, where the alpha pattern is dominated by a focal posterior cluster and the beta pattern is more spatially extended.

## 13. Appendix D. Selection of the group components

Two requirements governed which components were described: reproducibility across the sample, and a resolvable change in the coupling of the component’s own network inside its window.

Reproducibility was assessed by refitting. The nine subjects were split at random into groups of four and five, with both sessions of a subject kept in the same half, and the reduction and the decomposition were repeated independently within each half. Each half’s components were then matched one-to-one against those of the full sample by the absolute correlation of their mixing weights over all pairs. The scale of that correlation is set by the matching itself: random unit directions drawn in each half’s own principal subspace and matched in the same way reach a median |*r*| of 0.38 and a 95th percentile of 0.48, so a component was counted as recovered when its match exceeded |*r*| = 0.7. Over 40 splits this gives 80 halves, and the recovery rate of a component is the fraction of them in which it was recovered. Since each subject falls in exactly one half of every split, a component expressed by a single participant cannot be recovered in more than half of the halves; components were therefore retained when the Wilson 95% lower bound on their recovery rate was at least 0.50.

The second condition concerns effect size rather than consistency. The coupling of a component’s network inside its own window, averaged over the nine subjects, was required to reach 0.8 of the pre-stimulus standard deviation; components that were reproducible but moved their own pairs by less than this were not described further. Both quantities are listed for every component in Tables 1 and 2.

For each retained component we tested whether the excursion in its window is locked to the stimulus or a chance fluctuation of an ongoing process. The coupling time course of the network was circularly shifted in time by an independent random amount in each recording, 2000 times, and the group mean in the window was recomputed for every surrogate; the reported *p* is the fraction of surrogates reaching the observed value. Shifting preserves the amplitude, the shape and the autocorrelation of each recording, including those with unusually large deflections, and destroys only the alignment of those deflections to the stimulus.

To quantify how much of a component’s structure lies on local connections independently of any time window, we computed *E_L_*, the mean absolute mixing weight over all local pairs with at least one endpoint among the component’s sources, divided by the mean absolute weight over all remaining local pairs. Because any set of sources differs somewhat from the rest of the cortex, *E_L_* was referenced to a null of 2000 random source sets of the same size as that component’s node set, treated in the same way; components with *E_L_* above the median of that null are described as also loading on local connections. This measure uses the mixing weights only and is therefore independent of the window in which the component was examined.

Independent component analysis renders the time courses of the components independent but places no constraint on their mixing weights, so the pairs selected for one network also carry a share of other components. In the beta band the first four components capture sharp transients confined to single recordings, and their maps overlap appreciably with those of the networks described (|*r*| up to 0.64). Where indicated, their contribution was removed by regressing each recording’s coupling onto the canonical time courses of all twelve components and subtracting the fitted contribution of the first four, in the units in which the regression was performed, before any normalization. Results are reported both with and without this subtraction. Figures 19 and 20 show the full output of the decomposition in each band: the network of every component, its group-mean time course, and its local load *E_L_* against the null of random source sets of the same size.

In the alpha band all twelve components were reproducible in independently refitted halves of the sample, with matched spatial maps correlating at |*r*| = 0.71–0.91 against a null median of 0.38. Applying both criteria retained four of them. IC8 was reproducible but displaced its own pairs by only 0.12 pre-stimulus SD; the remaining seven were not recovered more often than a component carried by a single participant could be, and the selection therefore rests on reproducibility for seven of the eight components it rejects and on effect size for one.

The beta band shows the opposite pattern. Six components are reproducible, but only IC5 and IC6 also displace their own pairs by at least 0.8 pre-stimulus SD inside their windows; the remaining four move them by 0.51 to 0.73. Of the six components that fail reproducibility, four are the single-recording transients discussed in Section 3.2.3, whose local load (*E_L_* between 1.59 and 2.77) is the highest in either band.

## 14. Appendix E. Distal networks after removal of the dominant desynchronization subspace

This appendix reports a deliberately unfavourable variant of the similar analysis, in which the sensor-space subspace carrying the movement-related power modulation is removed, and only the recordings whose desynchronization survives that removal are kept. The coupling is then estimated from what remains, so that any distal network recovered cannot be a by-product of the local power effect that has been projected out. The procedure and the networks it returns are described below.

To ensure that subsequent connectivity analyses were restricted to regimes of genuine local desynchronization, we applied a spatial filtering procedure based on Common Spatial Patterns (CSP). CSP filters were computed by contrasting sensor-space covariance matrices from pre-stimulus and post-stimulus intervals, thereby isolating spatial components that maximally differentiated baseline activity from movement-related responses.

Figure 21A summarizes the effect of CSP-based projection on band-limited power changes across individual recordings. For both the alpha/mu and beta bands, CSP components capturing the strongest movement-related power suppression were identified and removed by projecting the data onto the orthogonal complement of the corresponding subspace. Across recordings, this procedure led to a systematic attenuation of the overall magnitude of desynchronization. While an increase in residual power relative to the original signals was observed in all recordings after CSP-based projection, the degree of this increase varied substantially. In a subset of sessions, the projected signals exhibited a pronounced enhancement of residual oscillatory power, indicating the presence of strongly expressed local synchronization components that were not fully suppressed by the CSP-defined subspace. These sessions were excluded from analysis.

**Figure 21:**
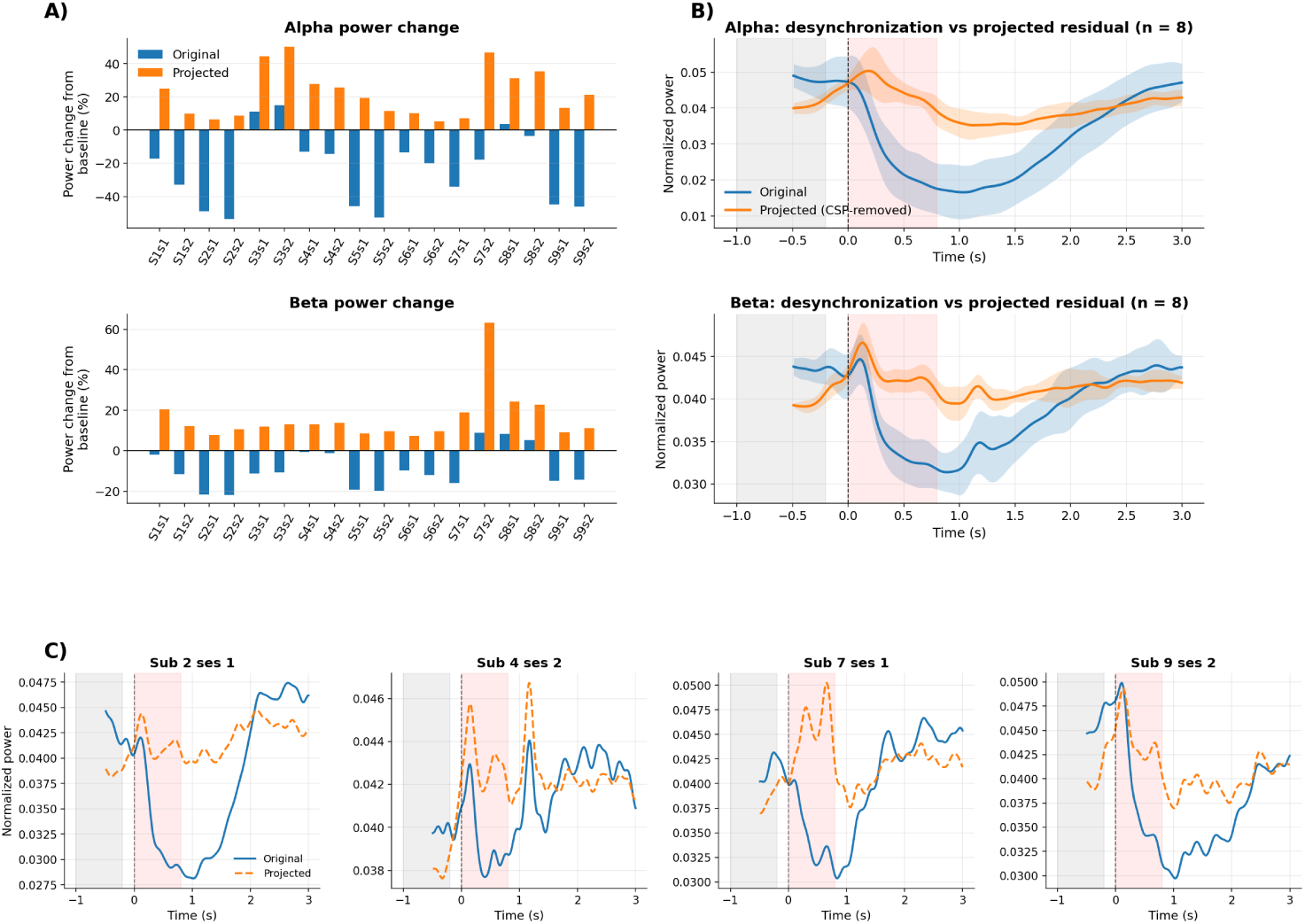
Effect of CSP-based spatial projection on movement-related desynchronization. (A) Percentage change in alpha/mu (top) and beta (bottom) band power from baseline to movement across individual recordings, shown before (original) and after projection onto the orthogonal complement of CSP components associated with maximal desynchronization. (B) Grand-average time courses of band-limited power in the alpha/mu (top) and beta (bottom) bands, shown before and after CSP-based projection. Averages are computed only over recordings selected for subsequent connectivity analysis. Shaded areas indicate variability across recordings. Vertical dashed lines denote stimulus and movement onset. (C) Representative single-subject examples of beta-band power time courses before and after CSP-based projection, illustrating the persistence of movement-related desynchronization at the individual recording level.

Panel B of Figure 21 shows the grand-average time courses of band-limited power before and after CSP-based projection, computed only over the recordings retained for subsequent analysis. In both frequency bands, movement-related power suppression remained clearly detectable after projection, despite a substantial reduction in amplitude relative to the original signals. This confirms that the selected recordings exhibited robust desynchronization.

Panel C of Figure 21 illustrates representative single-subject examples in the beta band, highlighting the heterogeneity of residual dynamics after CSP-based projection at the level of individual recordings. In some sessions (e.g., sub 2 ses 1 and sub 9 ses 2), a clear movement-related desynchronization remained visible after projection, with a temporal profile comparable to that observed in the original signals. In contrast, other sessions (e.g., sub 7 ses 1 and sub 4 ses 2) exhibited an enhancement of oscillatory power following projection, indicating a dominance of residual local synchronization in the projected signals.

Distal networks are recovered in both bands (Fig. 22). In the alpha band the 0.4–0.6 s window returns a bilateral pericentral network of 16 nodes, concentrated in postcentral and precentral cortex of both hemispheres. In the beta band the 1.2–1.4 s window returns a predominantly left-hemispheric parieto-frontal network of 17 nodes spanning inferior parietal, superior frontal and caudal middle frontal cortex. Inside their windows the distal coupling of these networks is at +0.24 and +0.92 prestimulus standard deviations, while the power of their own sources is at −4.79 and −2.52 and the coupling of the local pairs among their nodes at −1.96 and −0.47.

**Figure 22:**
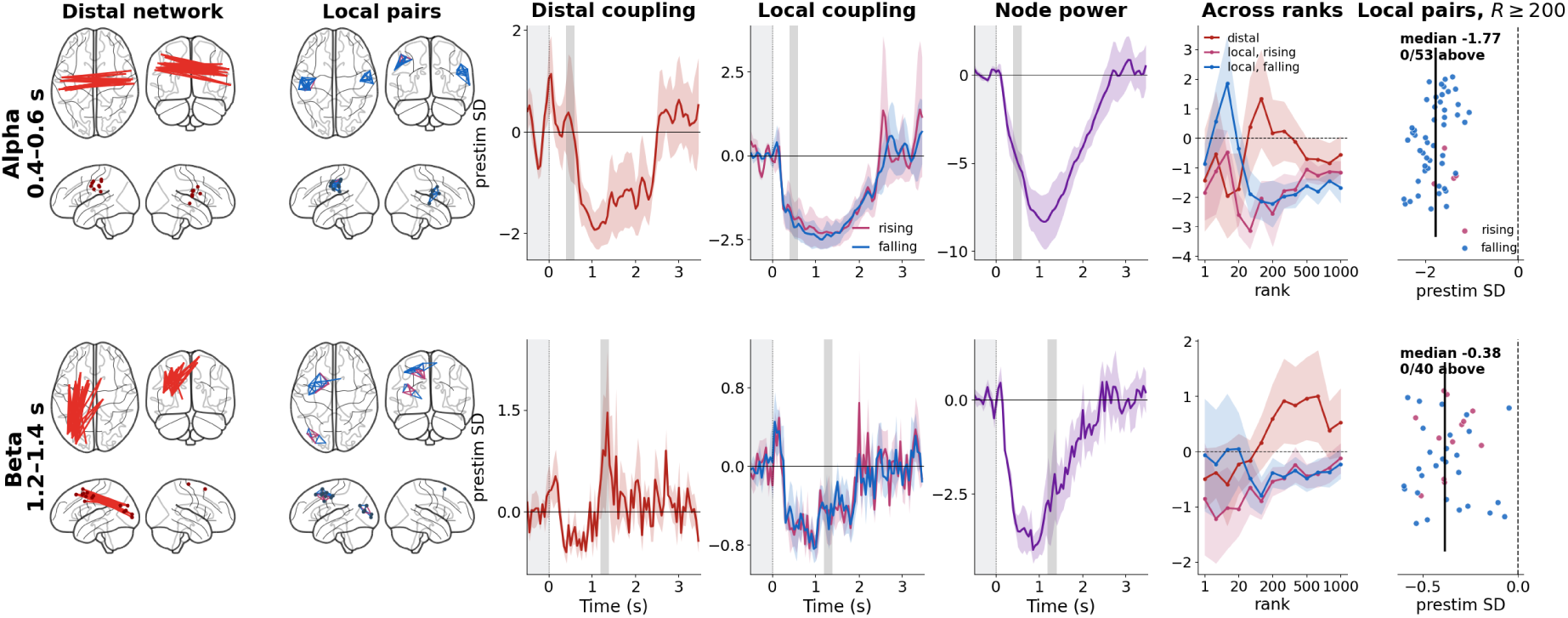
Distal networks recovered after removal of the dominant desynchronization subspace. Each row is one band and one window. From left to right: the thirty distal pairs with the strongest grand-averaged coupling increase in that window, in four views; the local pairs whose two endpoints are both sources of that network, coloured by whether their coupling rises (pink) or falls (blue) as the projection rank increases; the coupling of the distal network; the coupling of the two local subpopulations; the power of the participating sources; the window-mean coupling of the distal network and of the two local subpopulations as a function of the projection rank; and the level at which each local pair settles at high ranks (*R* ≥ 200), with the dashed line marking the prestimulus level, the black line the median over pairs, and the title giving that median and the number of pairs above the prestimulus level. Coupling is in units of each pair’s own prestimulus standard deviation and power in units of each source’s own prestimulus standard deviation; time courses are means over the four retained subjects ± SEM. Grey bands mark the prestimulus interval and the window.

The local pairs show no sign of accompanying synchronization at any projection rank. Of the 53 local pairs of the alpha network, 49 have coupling that falls as the rank increases, and of the 40 local pairs of the beta network, 28 do; at high ranks (*R* ≥ 200) the level at which they settle is −1.77 and −0.38 prestimulus standard deviations, and not a single local pair in either band lies above its own prestimulus level.

The rank profile of the distal estimate in the beta band is unambiguous. At the lowest ranks, when almost none of the power-dominated subspace has been removed, the distal coupling of the network sits *below* its prestimulus level (−0.49 at *R* = 1). It crosses zero near *R* = 100 and keeps growing to +0.92 at *R* = 300 and +1.00 at *R* = 600, while the local pairs of the same nodes remain below prestimulus at every rank. Spatial leakage behaves in exactly the opposite way: an apparent distal coupling produced by a shared local source is largest when the auto-topography subspace is left in and is progressively destroyed as that subspace is projected out. What is seen here is a distal interaction that only becomes visible once the power component has been removed, in a subsample from which the dominant desynchronization subspace had already been projected out at the sensor level.

In the alpha band the same profile is present but not monotone: the estimate rises from −1.42 at *R* = 1 to a maximum of +1.34 at *R* = 100, is +0.24 at the rank used throughout, and falls back below the prestimulus level at the highest ranks.

The subsample is too small for inference so all of these are descriptive statements, but the dissociation they describe is the one the main analysis reports, obtained here under conditions deliberately chosen to be unfavourable to it.

